# *In vivo* Characterization of the Critical Interaction between the RNA Exosome and the Essential RNA Helicase Mtr4 in *Saccharomyces cerevisiae*

**DOI:** 10.1101/2022.10.31.514520

**Authors:** Maria C. Sterrett, Daniela Farchi, Sarah E. Strassler, Lawrence H. Boise, Milo B. Fasken, Anita H. Corbett

**Affiliations:** Department of Biology, Emory University, Atlanta, Georgia; Biochemistry, Cell, and Developmental Biology Graduate Program, Emory University, Atlanta, Georgia; Department of Biochemistry, Emory University, Atlanta, Georgia; Department of Hematology and Medical Oncology, School of Medicine, Emory University, Atlanta, Georgia; Winship Cancer Institute, Emory University, Atlanta, Georgia

**Keywords:** RNA Exosome, Mtr4, EXOSC2, Rrp4, RNA processing, multiple myeloma, RNA helicase

## Abstract

The RNA exosome is a conserved molecular machine that processes/degrades numerous coding and non-coding RNAs. The 10-subunit complex is composed of three S1/KH cap subunits (human EXOSC2/3/1; yeast Rrp4/40/Csl4), a lower ring of six PH-like subunits (human EXOSC4/7/8/9/5/6; (yeast Rrp41/42/43/45/46/Mtr3), and a singular 3’-5’ exo/endonuclease DIS3/Rrp44. Recently, several disease-linked missense mutations have been identified in genes encoding the structural cap and core subunits of the RNA exosome. In this study, we characterize a rare multiple myeloma patient missense mutation that was identified in the cap subunit gene *EXOSC2*. This missense mutation results in a single amino acid substitution, p.Met40Thr, in a highly conserved domain of EXOSC2. Structural studies suggest this Met40 residue makes direct contact with the essential RNA helicase, MTR4, and may help stabilize the critical interaction between the RNA exosome complex and this cofactor. To assess this interaction *in vivo*, we utilized the *Saccharomyces cerevisiae* system and modeled the *EXOSC2* patient mutation into the orthologous yeast gene *RRP4*, generating the variant *rrp4 M68T*. The *rrp4 M68T* cells have accumulation of certain RNA exosome target RNAs and show sensitivity to drugs that impact RNA processing. Additionally, we identified robust negative genetic interactions the *rrp4 M68T* variant and RNA exosome cofactor mutants, particularly *mtr4* mutant variants. This study suggests that the *EXOC2* mutation identified in a multiple myeloma patient may impact the function of the RNA exosome and provides an *in vivo* assessment of a critical interface between the RNA exosome and Mtr4.

## INTRODUCTION

The RNA exosome is a highly conserved exo/endonuclease complex that has an essential role in 3’ to 5’ processing and degradation of nearly every species of RNA (Schneider and Tollervey 2013; Zinder and Lima 2017). First identified in *Saccharomyces cerevisiae* in a screen for ribosomal RNA processing (*rrp*) mutants (Mitchell *et al*. 1996; Mitchell *et al*. 1997), the RNA exosome is essential in all organisms studied thus far (Mitchell *et al*. 1997; Lorentzen *et al*. 2007; Hou *et al*. 2012; Lim *et al*. 2013; Pefanis *et al*. 2014). In addition to ribosomal RNA precursors, the RNA exosome processes a variety of small non-coding RNAs (ncRNAs), including small nuclear RNAs (snRNAs) and small nucleolar RNAs (snoRNAs) (Allmang *et al*. 1999; Van Hoof *et al*. 2000; Kilchert *et al*. 2016a; Fasken *et al*. 2020). The RNA exosome also plays roles in targeting RNA for degradation and decay, including non-functional or aberrant mRNAs and nuclear transcripts that result from pervasive transcription such as cryptic unstable transcripts (CUTs) in budding yeast or promoter upstream transcripts (PROMPTs) in humans (Wyers *et al*. 2005; Preker *et al*. 2008; Moore and Proudfoot 2009; Parker 2012; Schneider *et al*. 2012). The RNA exosome complex is composed of a 9-subunit structural core and a single exo/endonuclease [DIS3/DIS3L (human); Dis3/Rrp44 (budding yeast)]. As shown in Figure 1A, the 9-subunit structural core is composed of three S1/KH cap subunits (EXOSC1/2/3; Csl4/Rrp4/Rrp40) and a lower ring of six PH-like subunits (EXOSC4/5/6/7/8/9; Rrp41/Rrp46/Mtr3/Rrp42/Rrp43/Rrp45). The nuclear RNA exosome has an additional 3’-5’ exonuclease, EXOSC10/Rrp6, that associates with the complex and aids in nuclear RNA targeting and processing (Briggs *et al*. 1998; Wasmuth *et al*. 2014). Structural studies demonstrate that the overall organization of the RNA exosome is conserved (Figure 1B) suggesting not only evolutionary conservation of the RNA exosome function but structure as well (Makino *et al*. 2013; Schuller Jan *et al*. 2018; Weick *et al*. 2018). The vast array of targets and evolutionary conservation of the complex components indicates a fundamental role of the RNA exosome in several cellular processes, including but not limited to, maintaining genome integrity, translation, and cell differentiation through degradative and surveillance pathways (Ogami *et al*. 2018).

**Figure 1.**
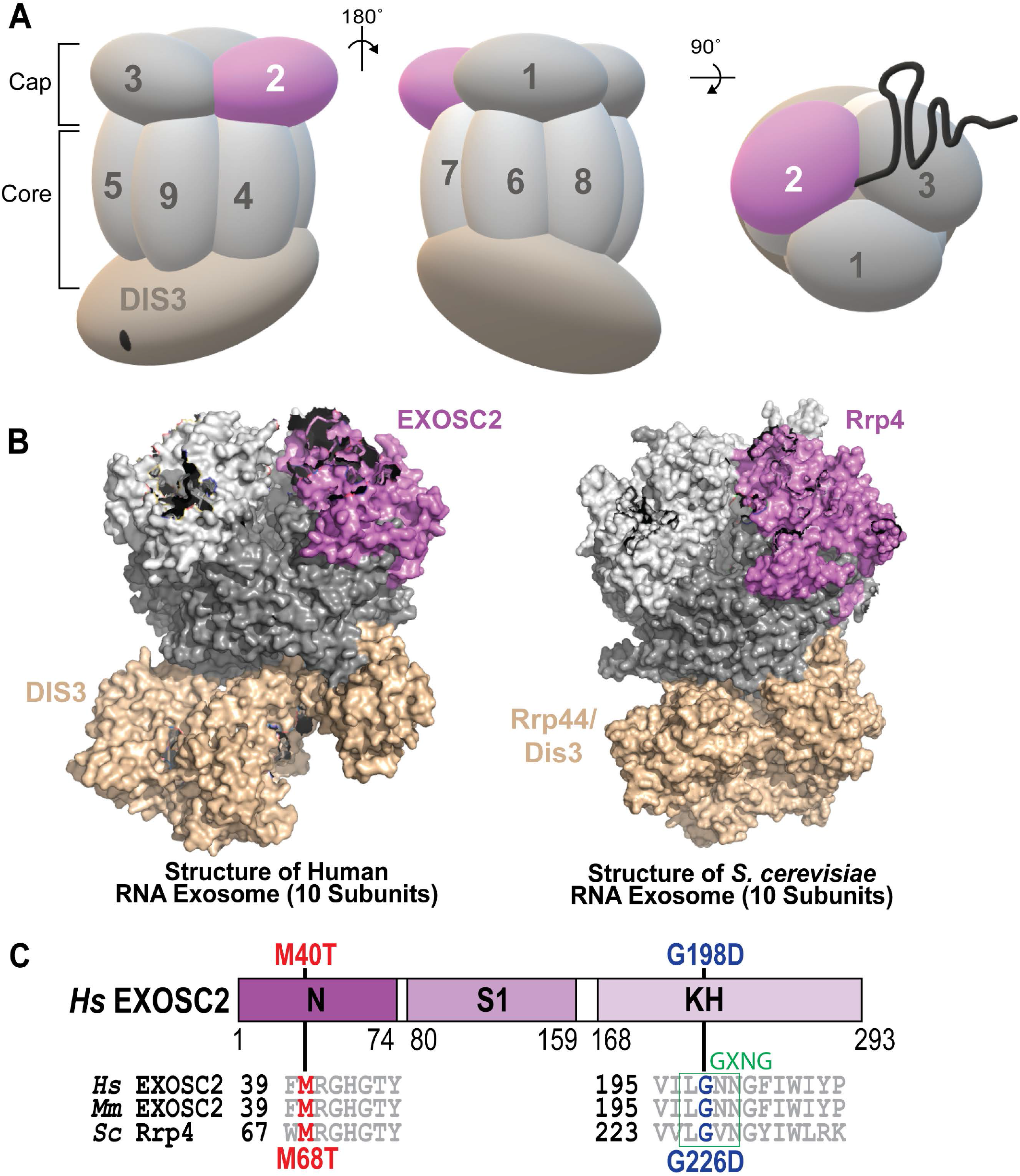
Overview of multiple myeloma linked amino acid substitutions in the human cap subunit EXOSC2 of the RNA exosome. (A) The RNA exosome is an evolutionary conserved ribonuclease complex composed of nine structural subunits (EXOSC1-9) and one catalytic subunit (DIS3) that form a “Cap” and “Core” ring-like structure. The 3-subunit cap at the top of the complex is composed of EXOSC1/Csl4 (Human/*S. cerevisiae*), EXOSC2/Rrp4, and EXOSC3/Rrp40 (labeled 1-3). The 6-subunit Core is composed of EXOSC4/Rrp41, EXOSC5/Rrp46, EXOSC6/Mtr3, EXOSC7/Rrp42, EXOSC8/Rrp43, and EXOSC9/Rrp45 (labeled 4-9). The DIS3/Dis3/Rrp44 catalytic subunit is located at the bottom of the complex. Together the cap and core form a barrel-like structure through which RNA is threaded to the catalytic DIS3/Dis3/Rrp44 subunit. Recent missense mutation in the gene encoding the EXOSC2 cap subunit (pink) have been identified in patients presenting with multiple myeloma. (B) Structural models of the human nuclear RNA exosome (left) [PDB 6D6R] (Weick *et al*. 2018) and the *S. cerevisiae* nuclear RNA exosome (right) [PDB 6FSZ] (Schuller *et al*. 2018) are depicted with the cap subunit EXOSC2/Rrp4 labeled and colored in pink. (C) Domain structure of EXOSC2/Rrp4. This cap subunit is composed of three domains: an N-terminal domain, a putative RNA binding S1 domain, and a C-terminal putative RNA binding KH (K homology) domain. A conserved “GxNG” motif identified in the KH domain is boxed in green (Oddone *et al*. 2007). The position of the disease-linked amino acid substitutions in human EXOSC2 are depicted above the domain structures. The amino acid substitution (p.Met40Thr) we report in a multiple myeloma patient is shown in red. An amino acid substitution (p.Gly198Asp) linked to SHRF is shown in blue (Di Donato *et al*. 2016). Sequence alignments of EXOSC2/Rrp4 orthologs from *Homo sapiens* (*Hs*), *Mus musculus* (*Mm*) and *S. cerevisiae* (*Sc*) below the domain structures show the highly conserved residues altered in disease in red and blue and the conserved sequences flanking these residues in gray.

RNA exosome specificity for a broad set of target transcripts is conferred in part through interactions with cofactor proteins, which aid the RNA exosome in target recognition, RNA unwinding, degradation, and catalysis in both the nucleus and the cytoplasm (Zinder and Lima 2017; Fasken *et al*. 2020). Many nuclear RNA exosome cofactors were first characterized in budding yeast, including the Rrp6 obligate binding partner Rrp47, Mpp6 and the essential 3’ to 5’ DExH box RNA helicase Mtr4 (De La Cruz *et al*. 1998; Mitchell *et al*. 2003; Lacava *et al*. 2005; Vaňáčová *et al*. 2005; Milligan *et al*. 2008), with orthologs now identified in the mammalian system (C1D, MPH6 and MTR4/MTREX) (Zinder and Lima 2017). Structural studies of the budding yeast and mammalian RNA exosome reveal that Rrp6/EXOSC10, Rrp47/C1D and Mpp6/MPH6 interact with the complex through conserved interfaces that form composite sites for interactions with other cofactors such as Mtr4/MTR4/MTREX (Schuch *et al*. 2014; Falk *et al*. 2017; Wasmuth *et al*. 2017; Schuller *et al*. 2018; Weick *et al*. 2018). The Mtr4 helicase assists in RNA substrate unwinding and plays a critical role in RNA exosome processing of the 5.8S rRNA precursor (7S rRNA) (De La Cruz *et al*. 1998; Taylor *et al*. 2014). Mtr4 also acts as part of larger complexes that aid the RNA exosome in nuclear RNA quality control, including the budding yeast TRAMP (Trf4/5-Air1/2-Mtr4 Polyadenylation) complex and the mammalian NEXT (Nuclear Exosome Targeting) complex (Houseley and Tollervey 2008; Houseley and Tollervey 2009; Weir *et al*. 2010; Lubas *et al*. 2011; Stuparevic *et al*. 2013; Falk *et al*. 2014; Kilchert *et al*. 2016a). Several studies that have dissected the role of Mtr4 in aiding the RNA exosome were performed in the *Saccharomyces cerevisiae* system, establishing a number of *mtr4* mutations that disrupt specific interactions and functions of the helicase (Kadowaki *et al*. 1994; Kadowaki *et al*. 1995; Liang *et al*. 1996; Weir *et al*. 2010; Taylor *et al*. 2014). Thus, genetic model systems are a tractable system to investigate interactions with these nuclear cofactors that impact RNA exosome function and studies in such systems can expand our understanding of the influence the RNA exosome can exert over various cellular processes and pathways (Cervelli and Galli 2021).

Given the variety of RNA exosome target RNAs and their link to many cellular processes, connections between the RNA exosome and human disease are not surprising. Many different human disease-linked mutations have been identified in genes encoding RNA exosome subunits (Fasken *et al*. 2020). Mutations in *DIS3*, which encodes the catalytic component of the RNA exosome in humans (Staals *et al*. 2010), are the fourth most common single nucleotide variation identified in multiple myeloma (∼10% of all newly diagnosed patients) (Chapman *et al*. 2011; Lohr *et al*. 2014). Multiple myeloma, which is a currently incurable cancer of the long-lived antibody-secreting plasma cells of the bone marrow, is the second most common hematologic malignancy accounting for 10-15% of incidence and 20% of deaths related to cancer of the blood and bone marrow (Alexander *et al*. 2007; Laubach *et al*. 2011). Multiple myeloma-associated *DIS3* mutations disrupt proper RNA degradation and processing in both mammalian cells and budding yeast mutant cells (Tomecki *et al*. 2014; Weissbach *et al*. 2015; Boyle *et al*. 2020). However additional mechanistic studies are required to understand how mutations in *DIS3*, and the function of the RNA exosome, could contribute to pathogenesis in multiple myeloma.

Human disease mutations have also been identified in the genes encoding the non-catalytic, structural subunits of the RNA exosome. Clinical studies have linked mutations in *EXOSC* genes to various, tissue-specific human pathologies comprising a growing family of diseases termed “RNA exosomopathies” (Wan *et al*. 2012; Boczonadi *et al*. 2014; Eggens *et al*. 2014; Di Donato *et al*. 2016; Burns *et al*. 2017; Slavotinek *et al*. 2020; Somashekar *et al*. 2021). RNA exosomopathy mutations have been found in all three genes that encode the cap subunits (*EXOSC1/2/3)* and several ring subunit genes (*EXOSC5/8/9*), with most being missense mutations that result in single amino acid substitutions in highly conserved domains of the subunits. Most RNA exosomopathy diseases are neurological, with mutations in *EXOSC1, EXOSC3, EXOSC5, EXOSC8*, and *EXOSC9* causing forms of cerebellar atrophy/degeneration and neuronopathies (Wan *et al*. 2012; Boczonadi *et al*. 2014; Eggens *et al*. 2014; Burns *et al*. 2018; Slavotinek *et al*. 2020; Somashekar *et al*. 2021). In contrast, patients with RNA exosomopathy mutations in *EXOSC2* have a complex syndrome known as SHRF that is characterized by <u>s</u>hort stature, <u>he</u>aring loss, retinitis pigmentosa and distinctive facies (OMIM #617763) (Di Donato *et al*. 2016). *In vivo* studies characterizing some of these *EXOSC* RNA exosomopathy mutations in *Saccharomyces cerevisiae* and *Drosophila melanogaster* suggest these pathogenic substitutions could differentially impact the function of the RNA exosome complex potentially through changes in RNA targeting and cofactor interactions (Fasken *et al*. 2017; Gillespie *et al*. 2017; Yang *et al*. 2019; De Amorim 2020; Morton *et al*. 2020; Slavotinek *et al*. 2020; Sterrett *et al*. 2021). Modeling these pathogenic amino acid substitutions in the budding yeast RNA exosome is an invaluable tool as several RNA exosomopathies have a small patient population, making analysis with patient tissue samples challenging. Therefore, by utilizing the budding yeast system, we can begin elucidating the functional and molecular consequences resulting from human disease mutations in RNA exosome genes can impact (Cervelli and Galli 2021).

In this study, we identify and characterize missense mutations in genes that encode the structural subunits of the human RNA exosome within multiple myeloma patients. We surveyed the ongoing longitudinal Multiple Myeloma Research Foundation (MMRF) study “Relating Clinical Outcomes in Multiple Myeloma to Personal Assessment of Genetic Profile” (CoMMpass) [ClinicalTrials.gov Identifier NCT01454297] to identify mutations in structural RNA exosome genes within multiple myeloma patients (Barwick *et al*. 2019). We focused on characterizing *EXOSC2 M40T*, a missense mutation that encodes an amino acid substitution EXOSC2 p.Met40Thr (M40T) in a highly conserved region of this cap subunit that interacts with the RNA helicase MTR4. To assess the effects of this amino acid substitution in EXOSC2 on the function of the RNA exosome, we utilized the budding yeast model system and generated a variant of the *S. cerevisiae* EXOSC2 ortholog, Rpr4, which models the patient EXOSC2 M40T substitution, Rrp4 M68T. As a comparative control within our studies, we included the Rrp4 G226D variant which models a SHRF-linked pathogenic amino acid substitution in EXOSC2 p.Gly198Asp (Sterrett *et al*. 2021). The *rrp4-G226D* cells, corresponding to the SHRF *EXOSC2* exosomopathy mutation, have defects in RNA exosome function, and are the only other budding yeast model of a disease-linked *EXOSC2* mutation (Sterrett *et al*. 2021). Our results show that the *rrp4-M68T* gene variant can replace the function of the essential *RRP4* gene. The *rrp4-M68T* and *rrp4-G226D* mutants show similar increases in specific RNA exosome target transcripts, suggesting shared defects in RNA processing. However, the *rrp4-M68T* mutant exhibits distinct negative genetic interactions with RNA exosome cofactor mutants, particularly *mtr4* mutants. Combined, our results suggest that the Rrp4 M68T amino acid substitution, which models the multiple myeloma associated substitution EXOSC2 M40T, alters RNA exosome function potentially by impacting the essential interaction between the complex and Mtr4. These data are the first *in vivo* characterization of this isolated multiple myeloma-associated mutation and give insight into the critical and conserved interactions between the RNA exosome and its cofactors.

## MATERIALS AND METHODS

### Media and Chemicals

All media were prepared by standard procedures (Adams *et al*. 1997). Unless stated otherwise, all chemicals were acquired from Fisher Scientific (Pittsburgh, PA), Sigma-Aldrich (St. Louis, MO), or United States Biological (Swampscott, MA).

### In silico protein structure predictions

The mCSM-PPI2 platform (Rodrigues *et al*. 2019), and the PyMOL viewer (The PyMOL Molecular Graphics System, Version 2.0 Schrödinger, LLC) (PyMOL) were used for structural modeling. Platforms were used with the cryo-EM structure (PDB 6D6Q) of the human nuclear RNA exosome at 3.45Å resolution (Weick *et al*. 2018) and the X-Ray diffraction structure (PDB 6FSZ) of the budding yeast nuclear RNA exosome at 4.60Å (Schuller Jan *et al*. 2018). The ConSurf server (Ashkenazy *et al*. 2010; Celniker *et al*. 2013; Ashkenazy *et al*. 2016) was used to assess the evolutionary conservation of the structure of both EXOSC2 and Rrp4.

### *Saccharomyces cerevisiae* strains and plasmids

All DNA manipulations were performed according to standard procedures (Sambrook *et al*. 1989). *S. cerevisiae* strains and plasmids used in this study are listed in Table S1. The *rrp4Δ* (yAV1103), *rrp4∆ mpp6∆* (ACY2471), and *rrp4∆ rrp47∆* (ACY2474) strains have been previously described (Schaeffer *et al*. 2009; Losh 2018; Sterrett *et al*. 2021). The *RRP45-TAP* (ACY2789) strain was obtained from Horizon Discovery Biosciences Limited and was previously described (Ghaemmaghami *et al*. 2003). The *mtr4∆* (ACY2532) strain was constructed by deletion of the genomic *MTR4* ORF in a wild-type (W303) strain harboring a [*MTR4; RRP4; URA3*] (pAC3714) maintenance plasmid by homologous recombination using *MTR4-UTR natMX4*. This *mtr4∆* (ACY2532) strain was then used for consecutive deletion of the genomic *RRP4* ORF to generate the *rrp4∆mtr4∆* (ACY2536) strain as previously described (Sterrett *et al*. 2021).

Construction of the untagged *RRP4* and *rrp4-G226D* plasmids (pAC3656 and pAC3659) and the 2x-Myc tagged *RRP4* and *rrp4-G226D* plasmids (pAC3669 and pAC3672) that contain native 3’ UTRs was reported previously (Sterrett *et al*. 2021). The *rrp4-M68T LEU2 CEN6* (pAC4206) and *rrp4-M68T-2xMyc LEU2 CEN6* (pAC4207) plasmids were generated by site-directed mutagenesis of the *RRP4* (pAC3656) or *RRP4-2xMyc* (pAC3669) plasmids using oligonucleotides containing the M68T missense mutation (Fwd 5’GAAAATACGTACCGTGACCTCT**CGT**CCAGATAGGGTCATCAGTGACC 3’, Rev 5’GGTCACTGATGACCCTATCTGG**ACG**AGAGGTCACGGTACGTATTTTC 3’) and the QuikChange II Site-Directed Mutagenesis Kit (Agilent). The *mtr4-F7A-F10A* (pAC4099) plasmid was generated as described previously (Sterrett *et al*. 2021). Similarly, the other *mtr4* mutant plasmids were constructed by site-directed mutagenesis of the *MTR4 HIS CEN6* plasmid (pAC4096) with the QuikChange II Site-Directed Mutagenesis Kit (Agilent) and oligonucleotides containing the corresponding missense mutations. The oligonucleotides used to generate the *mtr4-1* plasmid (pAC4103) contain the C942Y missense mutation (Fwd 5’CAAGCAGCAGCATTATTATCA**TAC**TTTGCATTCCAAGAACGCTG 3’, Rev 5’CAGCGTTCTTGGAATGCAAA**GTA**TGATAATAATGCTGCTGCTTG 3’). The oligonucleotides used to generate the *mtr4-R349E-N352E* plasmid (pAC4100) contain the R349E and N352E missense mutations (Falk *et al*. 2014) (Fwd 5’ GGTTGACGAAAAAAGTACCTTC**GAA**GAGGAA**GAA**TTCCAAAAAGCAATGGCGTCC 3’, Rev 5’ GGACGCCATTGCTTTTTGGAA**TTC**TTCCTC**TTC**GAAGGTACTTTTTTCGTCAACC 3’); The oligonucleotides used to generate the *mtr4-R1030A* plasmid (pAC4104) contain the R1030A missense mutation (Taylor *et al*. 2014) (Fwd 5’CGTTGATCAGAATGTTCAAG**GCA**TTAGAGGAATTGGTGAAGG 3’, Rev 5’CCTTCACCAATTCCTCTAA**TGC**CTTGAACATTCTGATCAACG 3’) and the oligonucleotides used to generate the *mtr4-E1033W* plasmid (pAC4105) contain the E1033W missense mutation (Taylor *et al*. 2014) (Fwd 5’GAATGTTCAAGAGATTAGAG**TGG**TTGGTGAAGGAGCTGGTAGAC 3’, Rev 5’GTCTACCAGCTCCTTCACCAA**CCA**CTCTAATCTCTTGAACATTC). Plasmids were confirmed through DNA sequencing.

### *Saccharomyces cerevisiae* transformations and growth assays

All *S. cerevisiae* transformations were conducted following the standard Lithium Acetate (LiOAc) protocol (Da *et al*. 2000). Strains were grown in liquid YEPD (1% yeast extract, 2% peptone, 2% dextrose, in distilled water) in a rotating shaker at 30°C overnight to saturation. Cultures were normalized to a concentration of OD600 = 0.33 in 10 mL YEPD, then incubated at 30°C for 3-8 hours depending on the severity of their growth defect. Cells were washed and resuspended to a concentration of 2 × 10^9^ cells/mL using TE/LiOAc. Single-stranded carrier DNA (5 μL; 10 mg/mL), PEG/TE/LiOAc (300 μL), and depending on reaction purpose, desired PCR product DNA or plasmid DNA, were added to cells. The mixture was incubated at 30°C in a shaker for 30 minutes. Following this incubation, DMSO (35 μL) was added and the cells were heat shocked for 15 minutes at 42°C, washed, and plated onto selective media.

Standard plasmid shuffle assays were performed to assess the *in vivo* function of the *rrp4* variants as well as genetic interaction with RNA exosome cofactor mutants. The *rrp4∆* (yAV1103) cells containing a *RRP4 URA3* maintenance plasmid and transformed with vector (pRS315) and transformed with *RRP4* (pAC3656), *rrp4-G226D* (pAC3659), *rrp4-M68T* (pAC4206), *RRP4-2xMyc* (pAC3669) or *rrp4-M68T-2xMyc* (pAC4207) plasmid were grown on Ura^-^ Leu^-^ minimal media control plates, which select for cells that contain both the *RRP4 URA3* maintenance plasmid as well as the *RRP4*/*rrp4 LEU2* plasmid, and 5-FOA Leu^-^ minimal media plates, which select for cells that lack the *RRP4 URA3* maintenance plasmid and contain only the *RRP4*/*rrp4 LEU2* plasmid. The plates were incubated at 30°C for 2-3 days and single colonies from the 5-FOA Leu^-^ minimal media plates were selected in quadruplicate and streaked onto selective Leu^-^ minimal media plates. The cells containing only the *RRP4/rrp4 LEU2* plasmid are referred to as *RRP4, rrp4-G226D* or *rrp4-M68T* cells. A similar strategy was used to generate *mtr4∆* (ACY2532) cells that contain only the *MTR4* (pAC4096) or *mtr4-1* (pAC4103) HIS3, CEN6 plasmid. The *mtr4∆* cells transformed with *MTR4* or *mtr4-1* were grown overnight and serially diluted and spotted onto Ura^-^ His^-^ minimal media plates and 5-FOA minimal media plates, which select for cells that lack the *URA3* maintenance plasmid and contain only the *MTR4*/*mtr4 HIS3* plasmid. Single colonies of cells containing only *MTR4* or *mtr4-1 HIS3* plasmid were collected in quadruplicate and are referred to as *MTR4* or as *mtr4-1* cells.

The *in vivo* function of the *rrp4-M68T* variant was assessed in growth assays on solid media and in liquid culture. For growth on solid media, *rrp4∆* (yAV1103) cells containing only *RRP4* (pAC3656), *rrp4-G226D* (pAC3659) or *rrp4-M68T* (pAC4206) were grown in 2 mL Leu^-^ minimal media overnight at 30°C to saturation. Cell concentrations were normalized to an OD_600_ = 0.5, and samples were serially diluted in 10-fold dilutions and spotted onto Leu^-^ minimal media plates, Leu^-^ minimal media plates containing 25 µM fluorouracil (5-FU), YEPD plates or YEPD plates containing 3% formamide, 150 mM hydroxyurea or 5 µg/ml phleomycin. Plates were grown at 25°C, 30°C, and 37°C for 2-3 days. For growth in liquid culture, cells were grown in 2 mL Leu^-^ minimal media overnight at 30°C to saturation, diluted to an OD_600_ = 0.01 in Leu^-^ minimal media in a 24-well plate, and growth at 37°C was monitored and recorded at OD_600_ in a BioTek® SynergyMx microplate reader with Gen5™ v2.04 software over 24 hr. For the results shown, each sample was performed in at least 3 independent biological replicates with 3 technical replicates for each biological sample. Doubling times were calculated using GraphPad Prism version 9.3.1 for Windows (www.graphpad.com), GraphPad Software, San Diego, California USA.

### Immunoblotting

To analyze protein expression levels of C-terminally Myc-tagged Rrp4 and Rrp4 M68T, *rrp4∆* (yAV1103) cells expressing only Rrp4-2xMyc (pAC3669) or Rrp4-M68T-2xMyc (pAC4207) were incubated in 2 mL of Leu-minimal medium at 30°C and grown to saturation overnight. The 10 mL cultures with an OD_600_ = 0.2 were prepared and incubated at 30°C or 37°C for 5 hr. Yeast cell pellets were collected by centrifugation and transferred to 2 mL screw-cap tubes. Cell pellets were flash frozen with liquid nitrogen and stored at -20°C. Yeast cell lysate was prepared by resuspending pellets in 0.3 mL of RIPA-2 Buffer (50 mM Tris-HCl, pH 8; 150 mM NaCl; 0.5% sodium deoxycholate; 1% NP40; 0.1% SDS) supplemented with protease inhibitors [1 mM PMSF; 3 ng/ml PLAC (pepstatin A, leupeptin, aprotinin, and chymostatin)], followed by addition of 300 µl glass beads. Lysates were placed in a Mini Bead Beater 16 Cell Disrupter (Biospec) for 6 × 1 min at 25°C with ice submersion intervals of 1 minute between rounds, and then centrifuged at 4°C at 12,000 RPM for 10 min. Protein lysate concentration was determined by Pierce BCA Protein Assay Kit (Life Technologies). Whole cell lysate protein samples (40 µg) in reducing sample buffer (50 mM Tris HCl, pH 6.8; 100 mM DTT; 2% SDS; 0.1% Bromophenol Blue; 10% Glycerol) were resolved on Criterion 4–20% gradient denaturing gels (Bio-Rad), transferred to nitrocellulose membranes (Bio-Rad) and Myc-tagged Rrp4 proteins were detected with anti-Myc monoclonal antibody 9B11 (1:2000; Cell Signaling). The 3-phosphoglycerate kinase (Pgk1) protein was detected using anti-Pgk1 monoclonal antibody (1:30,000; Invitrogen) as a loading control. For quantitation, ImageJ v1.4 software (National Institutes of Health, MD; http://rsb.info.nih.gov/ij/) was used to measure protein band areas and intensities. Protein percentages relative to Pgk1 were calculated using GraphPad Prismversion 9.3.1 for Windows (www.graphpad.com), GraphPad Software, San Diego, California USA.

### Co-Immunoprecipitations

To assess association of Rrp4 M68T with the RNA exosome complex, we utilized *RRP45-TAP* (ACY2789) cells expressing *RRP4-2xMyc* (pAC3669), *rrp4-G226D-2xMyc* (pAC3672) or *rrp4-M68T-2xMyc* (pAc4207) and immunoprecipitated Rrp45-TAP using the IgG Sepharose beads as previously described (Sterrett *et al*. 2021). Briefly, cell samples were grown in 2 mL Leu^-^ minimal media overnight at 30°C to saturation and 10-20 mL cultures with an OD_600_ = 0.2 were prepared and grown at 30°C for 5 hr. Yeast cell lysates were prepared by resuspending cell pellets in 0.5-0.75 mL of IPP150 Buffer (10mM Tris-HCl, pH 8; 150 mM NaCl; 0.1% NP40, PMSF) supplemented with protease inhibitors [1 mM PMSF; Pierce™ Protease Inhibitors (Thermo Fisher Scientific)], and 300 µL of glass beads. Cells were disrupted in a Mini Bead Beater 16 Cell Disrupter (Biospec) for 4-5 × 1 min at 25°C with 1 min on ice between repetitions. Crude lysate was transferred to a chilled microcentrifuge tube and remaining beads were washed with an additional 150 µL of IPP150 Buffer. Lysate was then cleared by centrifugation at 16,000 × *g* for 10 min at 4°C. Protein lysate concentration was determined by Pierce BCA Protein Assay Kit (Life Technologies). For input samples, 40 µg of cleared lysate was collected and frozen at -20°C. For co-immunoprecipitations, 1 mg of cleared lysate in IPP150 Buffer was prepared, 15-30 µL of a 1:1 bead slurry of either Pierce™ Anti-c-Myc Magnetic Beads (ThermoFisher) or IgG Sepharose® 6 Fast Flow Beads (GE Healthcare) was added, and samples were incubated at 4°C overnight with mixing. After overnight incubation, beads were washed three times in 1 mL IPP150 Buffer for 15 sec each (anti-Myc beads) or 5 min each (IgG Sepharose beads). Whole cell lysate input samples (40 µg) and total bound samples in reducing sample buffer were boiled for 5 min® at 100°C, resolved on 4–20% Criterion™ TGX precast polyacrylamide gels (Bio-Rad), transferred to nitrocellulose membranes (Bio-Rad). Levels of associated Rrp4-Myc proteins with the Rrp45-TAP tagged subunit were detected by immunoblotting. Myc-tagged Rrp4 proteins were detected with mouse anti-Myc monoclonal antibody 9B11 (1:2000; Cell Signaling). TAP-tagged Rrp45 protein was detected with peroxidase anti-peroxidase (PAP) soluble complex antibody produced in rabbit (1:5000, Sigma-Aldrich). The 3-phosphoglycerate kinase (Pgk1) protein was detected using anti-Pgk1 monoclonal antibody (1:30,000; Invitrogen) as a loading control.

### Genetic Interaction Analysis

To test genetic interactions between *rrp4-M68T* and RNA exosome cofactor/subunit deletion mutants, *rrp4∆ mpp6∆* (ACY2471), and *rrp4∆ rrp47∆* (ACY2474) cells containing only *RRP4* (pAC3656), *rrp4-G226D* (pAC3659) or *rrp4-M68T* (pAC4206) were grown in 2 mL Leu^-^ minimal media overnight at 30°C to saturation, serially diluted, and spotted on Leu^-^ minimal media plates. The plates were incubated at 30°C or 37°C for 3 days. Cells were also grown in liquid culture as described in *S. cerevisiae* transformation and growth assays method. The *rrp4∆ mpp6∆* (ACY2471) cells containing only *RRP4* (pAC3656), *rrp4-G226D* (pAC3659) or *rrp4-M68T* (pAC4206) were further assayed by being serially spotted onto Leu^-^ minimal media plates containing 25 µM 5-FU, YEPD plates or YEPD plates containing 3% formamide.

To test for genetic interactions between *rrp4-M68T* and *mtr4* mutants, *mtr4-F7A-F10A, mtr4-1, mtr4-R1030A*, and *mtr4-E1033W, rrp4∆ mtr4∆* (ACY2536) cells containing the [*MTR4; RRP4; URA3*] (pAC3714) maintenance plasmid were transformed with *RRP4* (pAC3656), *rrp4-G226D* (pAC3659) or *rrp4-M68T* (pAC4206) *LEU2* plasmid and selected on Ura^-^Leu^-^ minimal media plates. Transformed cells containing both the *URA3* maintenance plasmid and the *RRP4*/*rrp4* variant plasmid were subsequently transformed with *MTR4* (pAC4096), *mtr4-F7A-F10A* (pAC4099), *mtr4-1* (pAC4103), *mtr4-R1030A* (pAC4104), or *mtr4-E1033W* (pAC4105) *HIS3* plasmid and selected on Ura^-^Leu^-^His^-^ minimal media plates. The transformed cells were then streaked to onto 5-FOA Leu^-^ His^-^ plates to select for cells that did not contain the *URA3* maintenance plasmid. The resulting *rrp4∆ mtr4∆* cells containing only *RRP4, rrp4-G226D* or *rrp4-M68T LEU2* plasmid and *MTR4* or *mtr4* variant *HIS3* plasmid were grown in 2 mL Leu^-^ His^-^ minimal media overnight at 30°C to saturation, serially diluted, and spotted on Leu^-^ His^-^ minimal media plates. The plates were incubated at 30°C and 37°C for 3 days.

Cell growth was quantified on a scale from 0 to 5 across triplicate assays, with a score of “0” representing lethality and a score of “5” representing full growth across dilutions. Scores were averaged and displayed as a heatmap using GraphPad Prism version 9.3.1 for Windows (www.graphpad.com), GraphPad Software, San Diego, California USA.

### Total RNA isolatiosn

Total RNA from *RRP4, rrp4-G226D, rrp4-M68T, MTR4* or *mtr4-1* cells was isolated using the MasterPure™ Yeast RNA Purification Kit (Epicentre, Lucigen). Cells were incubated in 2 mL of Leu-minimal medium at 30°C and grown to saturation overnight. Cultures were diluted in 10 mL to an OD_600_ = 0.2 and further incubated at 37°C for 5 hours. Cells were pelleted by centrifugation, transferred to RNAse-free microcentrifuge tubes and flash frozen with liquid nitrogen. Frozen cell pellets were stored at -80°C. RNA isolation was performed according to the MasterPure™ Yeast RNA Purification Kit (Epicentre, Lucigen) manufacturer’s protocol. Total RNA was resuspended in 50 µL DEPC-treated water and stored at −80°C.

### qRT-PCR

All oligonucleotides used in this study are shown in Table S2. For analysis of steady-state RNA levels using quantitative PCR, three independent biological replicates of *RRP4, rrp4*-*G226D, rrp4-M68T, mtr4-1* and *MTR4* cells were grown in 2 mL Leu^-^ or His^-^ minimal media overnight at 30°C. Cultures (10 mL) with an OD_600_ = 0.2 were prepared from the saturated cultures and cells were grown at 37°C for 5 hr. Total RNA was isolated from cell pellets as described and 1 μg of total RNA was reverse transcribed to first strand cDNA using the M-MLV Reverse Transcriptase (Invitrogen) according to manufacturer’s protocol. Quantitative PCR was performed on technical triplicates of cDNA (10 ng) from three independent biological replicates using gene specific primers (0.5 mM; Table S2), QuantiTect SYBR Green PCR master mix (Qiagen) on a StepOnePlus Real-Time PCR machine (Applied Biosystems; Tanneal=55°C, 44 cycles). *ALG9* or *PGK1* was used as an internal control. The mean RNA levels were calculated by the ΔΔCt method (Livak and Schmittgen 2001). Statistical analysis comparing the control cells (*RRP4* or *MTR4*) and the mutant cells (*rrp4* or *mtr4-1*) was performed by t-test (α<0.05) using GraphPad Prism version 9.3.1 for Windows (www.graphpad.com), GraphPad Software, San Diego, California USA.

## DATA AVAILABILITY STATEMENT

Strains and plasmids summarized in Table S1 and oligonucleotides are shown in Table S2. These reagents are available upon request. Genomic data from CoMMpass are available at dbGaP with the accession number phs000748.v7.p4. The authors affirm that all data necessary for confirming the conclusions of the article are present within the article, figures, and tables.

## RESULTS

### EXOSC2 p.Met40Thr substitution is located within a conserved region of the cap subunit that interacts with the essential helicase MTR4

Mutations in the gene *DIS3*, which encodes the catalytic component of the RNA exosome, are commonly found in patients diagnosed with multiple myeloma (Chapman *et al*. 2011; Lohr *et al*. 2014), suggesting a link between RNA exosome function and disease pathology. We therefore considered whether mutations in the other components of the RNA exosome would be found in multiple myeloma patients. Missense mutations in *EXOSC* genes, which encode the structural subunits of the RNA exosome, were identified in multiple myeloma patients through interrogating the ongoing longitudinal Multiple Myeloma Research Foundation (MMRF) study “Relating Clinical Outcomes in Multiple Myeloma to Personal Assessment of Genetic Profile” (CoMMpass) [ClinicalTrials.gov Identifier NCT01454297]. A total of 1,154 newly diagnosed multiple myeloma patients were enrolled in CoMMpass and profiled by genomic testing and tissue sampling throughout treatment. The molecular profiling collected through CoMMpass reveals several rare missense mutations within *EXOSC* genes. One patient missense mutation identified within exon 1 of *EXOSC2* encodes EXOSC2 p.Met40Thr (M40T), which is located in a highly conserved region of the N-terminal domain of EXOSC2 (Figure 1C). Notably, EXOSC2 Met40 lies within a key binding interface between the human RNA exosome and the RNA helicase MTR4 (Weick *et al*. 2018).

The patient with the *EXOSC2 M40T* mutation also had chromosomal aberrations including a chromosomal translocation t(11;14) and hyperdiploidy disease. The chromosomal translocation t(11;14) is an IgH translocation which is an initiating event that occurs frequently in multiple myeloma (∼15-20% of patients) (Bergsagel *et al*. 1997). From the CoMMpass dataset, we determined that the variant allele frequency is 0.2266, however the copy number of the chromosome 9 *EXOSC2* locus is 2.6, suggesting that this *EXOSC2* allele is found on the extra copy of ch9 that is present in over half the cells in the patient. Based on these findings, we conclude that this hyperdiploidy of chromosome 9 occurred after the t(11;14) translocation event and that the *EXOSC2* mutation either co-occurred with the chromosomal gain or shortly after.

To explore how EXOSC2 M40T could alter the function of the RNA exosome complex, we modeled the EXOSC2 M40T amino acid substitution using a recent structure of the human RNA exosome in complex with the essential RNA helicase MTR4 (Weick *et al*. 2018). MTR4 makes several direct contacts with the RNA exosome, forming a binding interface with a total surface area of 1,440Å (Weick *et al*. 2018). Among the direct contacts between MTR4 and the RNA exosome complex, the N-terminal domain of EXOSC2 interacts with MTR4 through an aliphatic surface that includes Met40. As shown in Figure 2A, EXOSC2 Met40 engages with a hydrophobic pocket of MTR4 including I1014. An amino acid substitution of Thr40, while unlikely to disrupt the aliphatic surface, could disrupt the hydrophobic interaction at this contact given the polar, shortened side chain of threonine (Figure 2A). The EXOSC2 M40T substitution could therefore destabilize the interface between the N-terminal domain of EXOSC2 and MTR4. We also modeled the amino acid substitution in the budding yeast EXOSC2 ortholog, Rrp4, using a recent structure of the *S. cerevisiae* RNA exosome (Schuller *et al*. 2018). As shown in Figure 2B, the budding yeast RNA exosome in complex with Mtr4 shows structural similarities to the human complex, with Rrp4 interacting directly with Mtr4. Rrp4 Met68, corresponding to the EXOSC2 Met40 residue, engages with the helicase directly through hydrophobic interactions at the binding interface.

**Figure 2.**
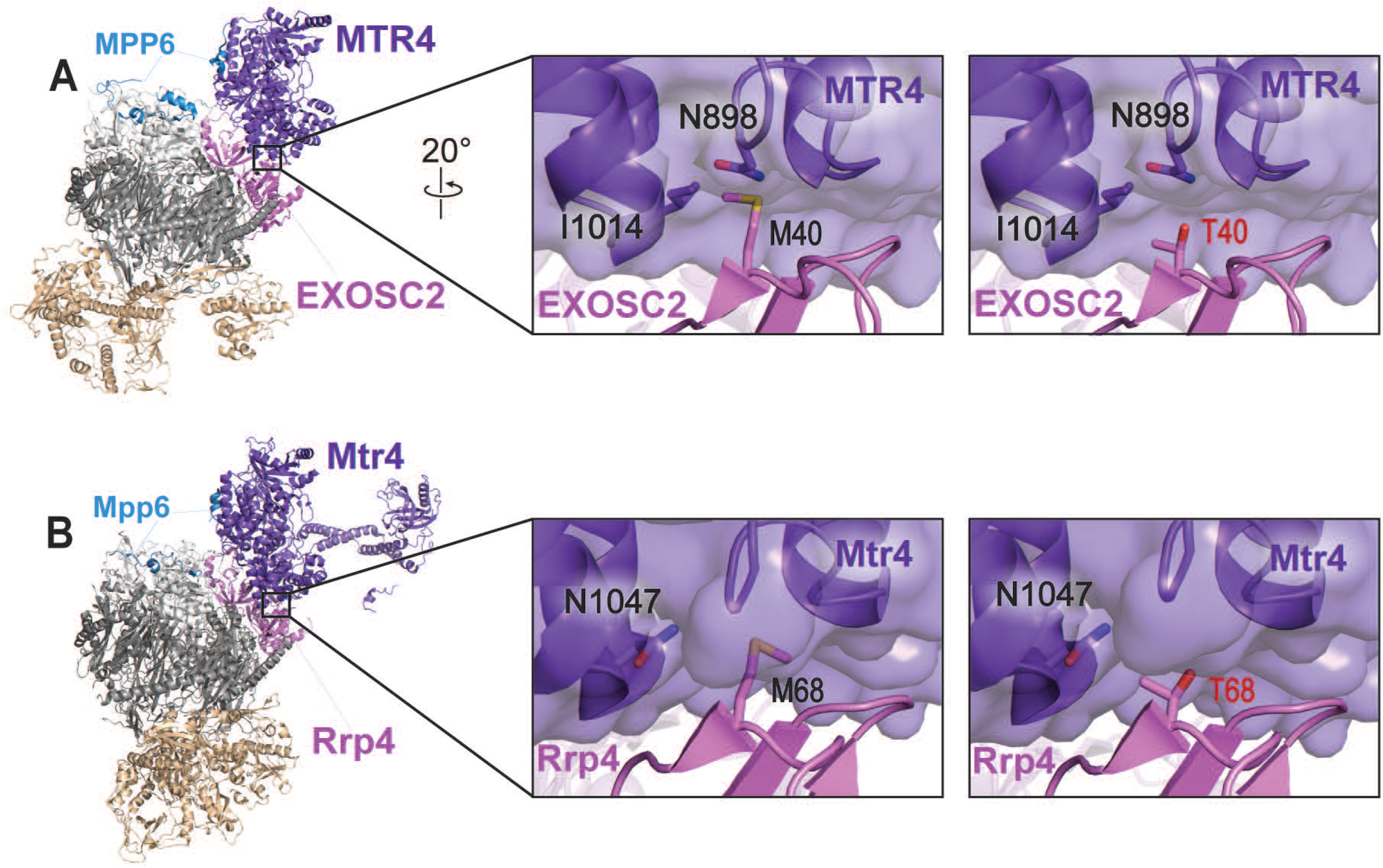
Modeling the multiple myeloma EXOSC2 M40T amino acid substitution in the human EXOSC2 cap subunit and the *S. cerevisiae* ortholog Rrp4. (A) Structural modeling of the human EXOSC2 p.Met40Thr (M40T) amino acid substitution identified in a patient with multiple myeloma (PDB 6D6R) (Weick *et al*. 2018). The full structure of the human RNA exosome with the associated cofactor MTR4 (purple) is depicted with a zoomed-in representation of the interface between EXOSC2 (pink) and MTR4. Modeling of the native EXOSC2 Met40 (M40) residue (left) or the multiple myeloma-associated EXOSC2 Thr40 (T40) residue (right) is shown. The EXOSC2 Met40 residue is located in the N-terminal domain of EXOSC2, within a conserved aliphatic interface with MTR4. EXOSC2 Met40 and MTR4 associate through hydrophobic interactions, which includes contacts with MTR4 Ile1014 (I1014). (B) Structural modeling of the budding yeast Rrp4 Met68Thr (M68T) amino acid change, corresponding to EXOSC2 p.Met40Thr, in the budding yeast RNA exosome (PDB 6FSZ) (Schuller *et al*. 2018). The full structure of the budding yeast RNA exosome complex with the associated MTR4 ortholog, Mtr4 (purple), is depicted on the left. A zoomed-in representation of the interface between Rrp4 (pink) and Mtr4 are shown, modeling the native Rrp4 Met68 (M68) residue (left) or the modeled multiple myeloma associated substitution Rrp4 Thr68 (T68) residue (right). The Rrp4 Met68 residue is conserved between human and yeast and is located in the N-terminal domain of Rrp4. Similar to EXOSC2 Met40, Rrp4 Met68, associates with the helicase Mtr4 through primarily hydrophobic interactions, including contacts with several glycine residues in a neighboring loop of Mtr4. Parts of the human nuclear cofactor protein MPP6/MPH6 (blue) and the budding yeast ortholog Mpp6 (blue) are also resolved in the structures shown in (A) and (B). Both MPP6/MPH6 and Mpp6 aid in stabilizing the interaction of the RNA helicase with the RNA exosome in addition to the direct interface made between EXOSC2 and MTR4 in humans or Rrp4 and Mtr4 in budding yeast (Falk *et al*. 2017; Wasmuth *et al*. 2017).

Introduction of Thr40 would most likely disrupt this contract, similar to our predictions for the M40T substitution in EXOSC2. Furthermore, the region surrounding Rrp4 Met68 is structurally synonymous to the aliphatic surface surrounding EXOSC2 Met40, allowing for us to assess the effects of the EXOSC2 M40T amino acid substitution at the conserved interface within the yeast system.

We further explored the conservation of the binding interface of human EXOSC2 as compared to budding yeast Rrp4 using the bioinformatics tool ConSurf (Figure S1). The ConSurf server estimates the evolutionary conservation of amino acids of a protein based on phylogenetic trees between homologous sequences, providing conservation rates for each residue that reflect both functional and structural importance (Ashkenazy *et al*. 2010; Celniker *et al*. 2013; Ashkenazy *et al*. 2016). Consistent with the sequence alignment (Figure 1C) and structural modeling, this tool predicts high conservation for both EXOSC2 Met40 and Rrp4 Met68 and surrounding residues (Figure S1). Additionally, ConSurf estimates high conservation rates at each site of contact between EXOSC2 and Rrp4 with the helicase MTR/Mtr4, further supporting the evolutionary importance of this interaction.

### *Saccharomyces cerevisiae rrp4-M68T* mutant cells that model the multiple myeloma *EXOSC2 M40T* variant show sensitivity on drugs that impact RNA metabolism

To assess the functional consequences of the EXOSC2 M40T amino acid substitution, we generated the corresponding amino change in the *S. cerevisiae* ortholog Rrp4, M68T (Figure 1C). We first performed a plasmid shuffle growth assay in which cells deleted for the genomic copy of *RRP4* are transformed with plasmids containing different *rrp4* alleles (See *Materials and Methods)*. This approach ensures that the background for all variants that are compared to one another is identical (Sikorski and Hieter 1989). This growth assay reveals the *rrp4-M68T* allele can replace the essential *RRP4* gene as the *rrp4-M68T* cells grow similarly to control cells expressing wild-type *RRP4* at all temperatures examined (Figure 3A). As a comparison, we included cells expressing the *rrp4-G226D* allele as the sole copy of the essential *RRP4* gene. The *rrp4-G226D* mutant allele models a known SHRF pathogenic amino acid change that has been shown to cause impaired RNA exosome function *in vivo* (Sterrett *et al*. 2021). As previously reported, cells expressing only *rrp4-G226D* show impaired growth at 37°C (Figure 3A). Furthermore, we assessed the growth of both the *rrp4-M68T* and *rrp4-G226D* mutant cells using a liquid growth assay and quantified doubling times (Figure 3B, 3C). These data confirm that the growth of *rrp4-M68T* cells does not differ significantly from wild-type *RRP4* cells.

**Figure 3.**
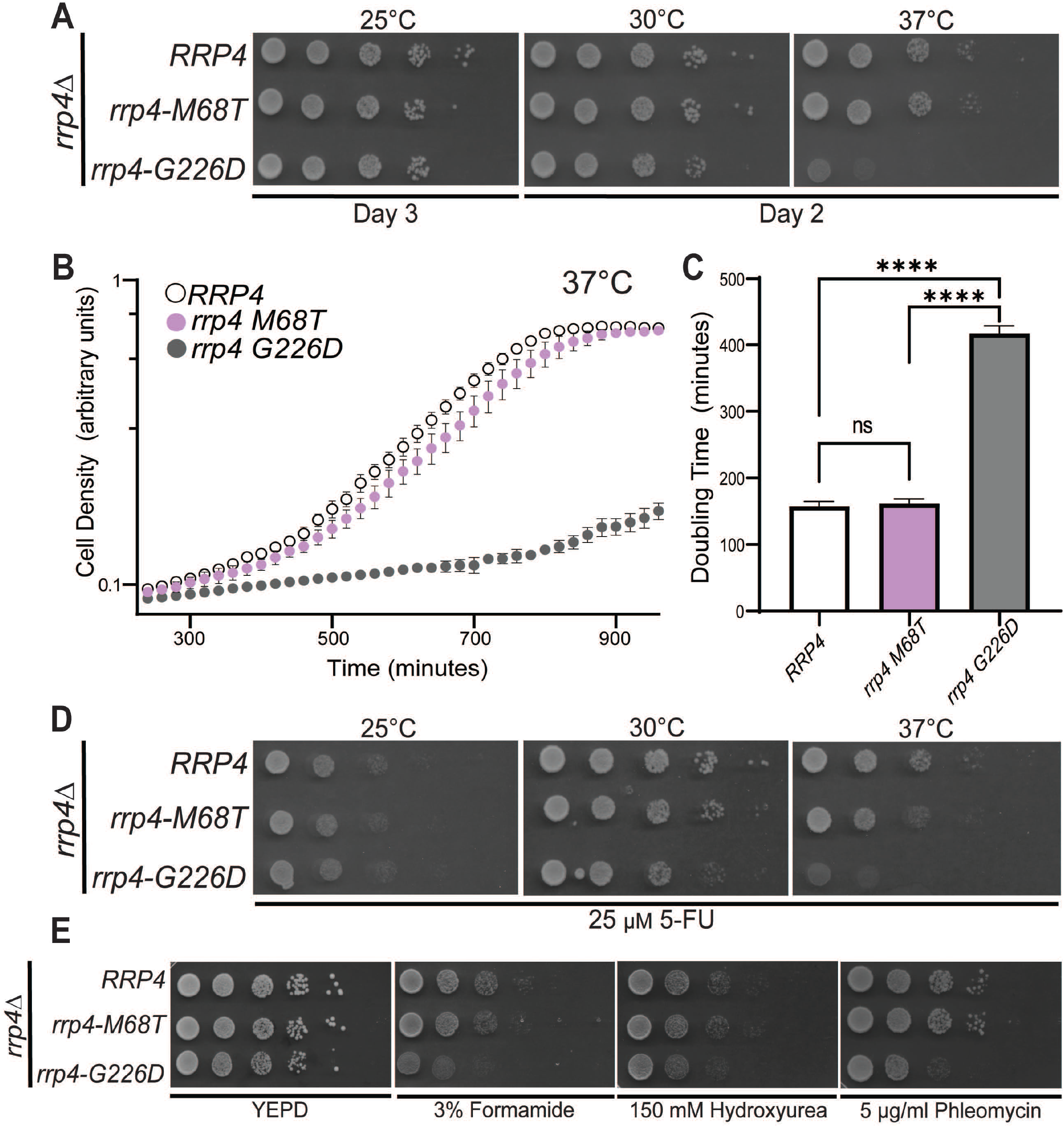
*S. cerevisiae rrp4-M68T* mutant cells that model the EXOSC2 M40T variant identified in multiple myeloma patients show impaired function on drugs that impact RNA processing. *S. cerevisiae* cells expressing Rrp4 variants that model the multiple myeloma amino acid change or, as a control, the previously characterized (Sterrett *et al*. 2021) SHRF-linked amino acid change found in EXOSC2 were generated as described in *Materials and Methods*. (A-B) The *rrp4∆* cells expressing only *RRP4* or mutant *rrp4* were serially diluted, spotted onto solid selective media grown at the indicated temperatures or grown in liquid media at 37°C with optical density measurement used to assess cell density over time. The doubling time of these cells grown in liquid media is quantified and graphed in (C). (D) The *rrp4∆* cells expressing either *RRP4* or *rrp4-M68T* were serially diluted, spotted onto solid selective media containing 25 µM fluorouracil (5-FU) and grown at the indicated temperatures. Images shown are after two days of growth. (E) The *rrp4∆* cells expressing only *RRP4* or *rrp4-M68T* were serially diluted and spotted onto solid YEPD media containing 3% formamide, 150 mM hydroxyurea or 5µg/ml phleomycin and grown at 30°C. Images shown are after two days of growth. In all assays performed, *rrp4-G226D* cells, previously reported to be severely impaired at 37°C, were included as a control (Sterrett *et al*. 2021). Data shown are representative of three independent experiments (n = 3).

To explore whether the *rrp4-M68T* mutation sensitizes cells to altered RNA processing, we tested for growth defects when cells are grown on media containing 5-fluorouracil (5-FU) (Giaever *et al*. 2004; Hoskins and Scott Butler 2007) (Figure 3D). The *rrp4-M68T* cells show a slight growth defect compared to wild-type *RRP4* cells at 30°C when grown on solid media containing 25 µM 5-FU. This growth defect is more evident when the cells are challenged with both 37°C and 25 µM 5-FU. As a comparison, the *rrp4-G226D* cells show a severe growth defect when grown on solid media containing 5-FU both at 30°C and 37°C. To further explore whether the *rrp4-M68T* cells exhibit other changes in cell growth, we tested for growth defects when cells are grown on media containing chemicals that disrupt different cellular pathways (Figure 3E). Formamide alters RNA metabolism (Hoyos-Manchado *et al*. 2017), hydroxyurea impairs DNA synthesis (Slater 1973), and phleomycin acts as a mutagen by introducing double-strand breaks in DNA (Suzuki *et al*. 1970). The *rrp4-M68T* cells do not show any increased sensitivity when grown at 30°C on solid media containing 3% formamide, 150 mM hydroxyurea or 5 µg/ml phleomycin (Figure 3E). In contrast, the *rrp4-G226D* cells show enhanced growth defects at 30°C when grown on solid media containing 3% formamide, 150 mM hydroxyurea or 5 µg/ml phleomycin. Taken together, these data suggest that the *rrp4-M68T* cells are sensitive to defects in RNA processing but do not exhibit the same extent of disrupted cellular pathways as the previously studied *rrp4-G226D* cells, which model a pathogenic RNA exosomopathy mutation that has severely impaired RNA exosome function *in vivo* (Sterrett *et al*. 2021).

### *rrp4-M68T* cells have impaired RNA exosome function in processing RNA targets linked to Mtr4-RNA exosome interactions

To further assess the *in vivo* consequences on RNA exosome function of the *rrp4-M68T* variant, we measured the steady-state levels of several RNA exosome targets in *rrp4-M68T* cells using RT-qPCR. We assessed the steady-state level of precursor RNAs that are targeted by the RNA exosome and are impacted by *mtr4* mutant alleles, including the telomerase component RNA *TLC1*, which is processed by the RNA exosome in a manner dependent on TRAMP complex association, and the 3’ extended forms of *U4* snRNA and *snR33* snoRNA (Van Hoof *et al*. 2000; Houseley *et al*. 2006; Coy *et al*. 2013). In this analysis, we included both *rrp4-G226D* and *mtr4-1* cells as comparative controls. The *mtr4-1* cells have a missense mutation in *MTR4* that results in accumulation of polyadenylated targets within the nucleus (Kadowaki *et al*. 1994; Kadowaki *et al*. 1995; Liang *et al*. 1996; Weir *et al*. 2010). We detect increases in the steady-state level of both mature and precursor *TLC1* in *rrp4-M68T* cells similar to that observed in *rrp4-G226D* cells (Figure 4A) (Sterrett *et al*. 2021). Both mature and precursor *TLC1* steady-state levels are significantly increased in *mtr4-1* cells.

**Figure 4.**
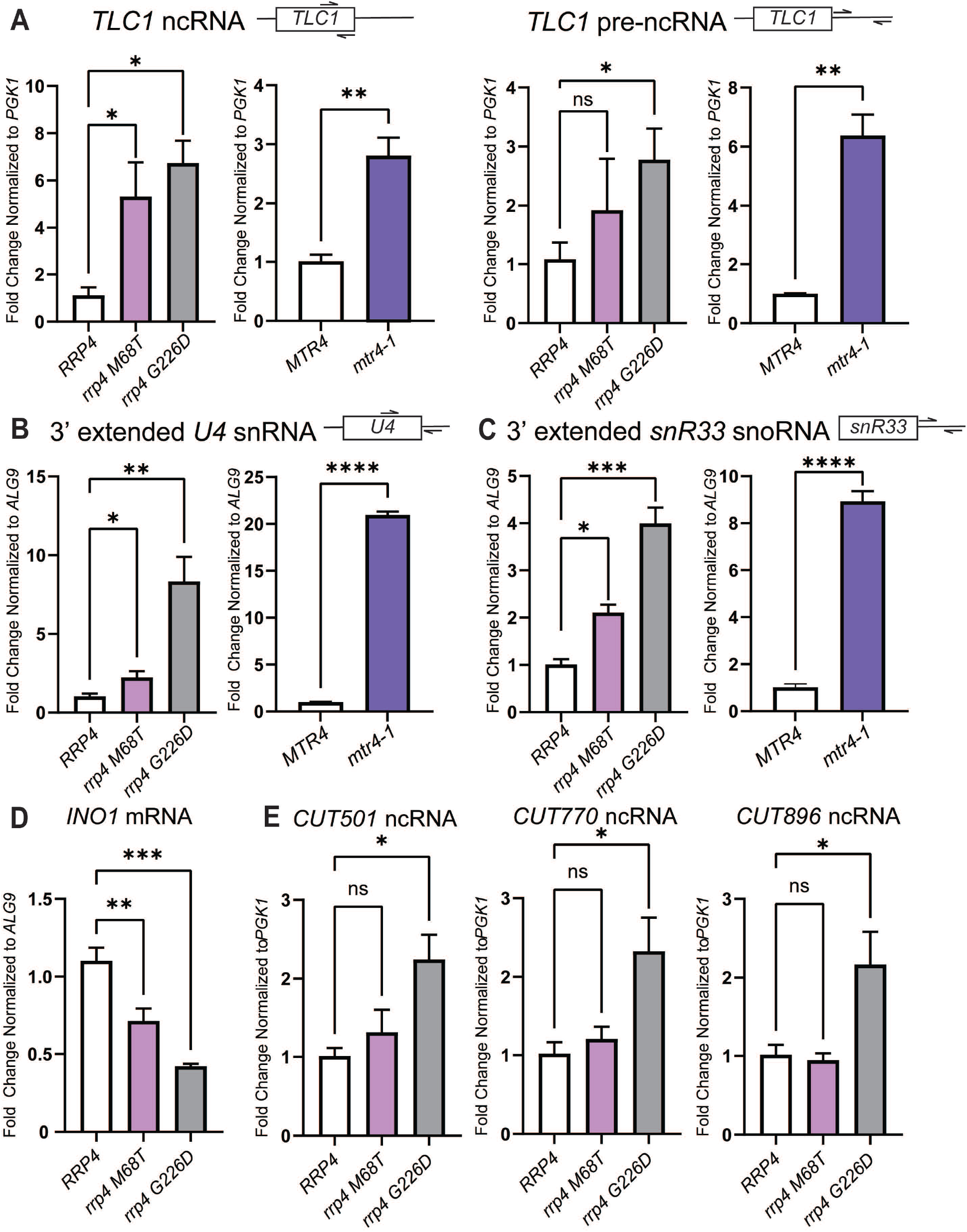
The *rrp4-M68T* mutant cells show elevated levels of specific RNA exosome target transcripts that depend on the Mtr4-RNA exosome interaction *in vivo*. The steady-state level of several RNA exosome target transcripts was assessed in *rrp4-M68T* cells (denoted in pink). The steady-state levels of these RNAs were also assessed in the previously described *rrp4* variant *rrp4-G226D* as a control (denoted in gray). (A) The *rrp4-M68T* cells show an elevated steady-state level of mature *TLC1* telomerase component ncRNA relative to *RRP4* cells. The steady-state level of the precursor *TLC1* ncRNA in *rrp4-M68T* cells follows this upward trend though not statistically significant compared to *RRP4* cells. This increase in both mature and precursor *TLC1* is also observed in *mtr4-1* (denoted in purple) compared to the wild-type control *MTR4*. (B) The *rrp4-M68T* cells show an elevated steady-state level of 3’-extended pre-*U4* snRNA relative to *RRP4* cells. The *rrp4-G226D* mutant cells and *mtr4-1* cells have an even higher steady-state levels of this pre-*U4* snRNA when compared to the *RRP4* and *MTR4* controls, respectively. (C) The *rrp4-M68T* cells show an elevated steady-state level of 3’ extended *snR33* snoRNA relative to *RRP4* cells that is similar to the increase observed in the *rrp4-G226D* mutant cells and *mtr4-1* cells. (D) The *rrp4-M68T* cells show a decreased steady-state level of the mRNA transcript *INO1* compared to wild-type *RRP4* control cells. A decrease in this mRNA was shown previously in *rrp4-G226D* cells (Sterrett *et al*. 2021). (E) The steady-state levels of non-coding, cryptic unstable transcripts, *CUT501, CUT770*, and *CUT896*, are not significantly increased in *rrp4-M68T* cells compared to control as shown previously in the *rrp4-G226D* mutant cells (Sterrett *et al*. 2021). In (A-E), total RNA was isolated from cells grown at 37°C and transcript levels were measured by RT-qPCR using gene specific primers and graphed as described in *Materials and Methods*. Gene specific primer sequences are summarized in Table S2. The location of primers specific to the ncRNA transcripts are graphically represented by the cartoons above each bar graph. Within the cartoon transcript, the box represents the body of the mature transcript. Error bars represent standard error of the mean from three biological replicates. Statistical significance of the RNA levels in *rrp4* variant cells relative to *RRP4* cells and in the *mtr4*-1 cells relative to *MTR4* cells is denoted by an asterisk (\**p*-value ≤ 0.05; \*\**p-*value ≤ 0.01, \*\*\**p-*value ≤ 0.001, \*\*\*\**p-*value ≤ 0.0001).

Furthermore, we detect a significant increase in the steady-state level of the 3’ extended forms of the *U4* snRNA and *snR33* snoRNA in the *rrp4-M68T* cells (Figure 4B and 4C). This increase in the levels of the extended form of these target RNAs is also observed in the *rrp4-G226D* cells and, to an even larger extent, the *mtr4-1* cells. We also assessed steady-state levels of 5.8S rRNA precursors in *rrp4-M68T* and found no accumulation compared to wild-type *RRP4* cells (Figure S2), although levels of both mature 5.8S rRNA and pre-5.8S rRNA do increase in *rrp4-G226D* cells which supports previous observations that the *rrp4-G226D* cells exhibit accumulation of 7S rRNA (Sterrett *et al*. 2021).

We also measured the steady-state level of select targets that are impacted within *rrp4-G226D* cells (Sterrett *et al*. 2021). We assessed the target *INO1*, which encodes inositol-3-phosphate synthetase (Donahue and Henry 1981; Klig and Henry 1984). *INO1* mRNA has previously been identified as a transcript bound to the catalytic subunit of the RNA exosome (Delan-Forino *et al*. 2017), and was the most significantly decreased transcript in a previous RNA-Seq analysis of the *rrp4-G226D* cells (Sterrett *et al*. 2021). In *rrp4-M68T* cells, the steady-state level of *INO1* is significantly decreased, similar to results for *rrp4-G226D* (Figure 4D). We also assessed three cryptic unstable transcripts (CUTs) that accumulate within *rrp4-G226D* cells (Sterrett *et al*. 2021). The steady-state levels of these three CUTs are not significantly increased in *rrp4-M68T* cells (Figure 4E). Taken together, these data suggest that the *rrp4-M68T* cells have some molecular consequences resulting from the modeled multiple myeloma amino acid substitution; however, they differ from those resulting from the modeled SHRF substitution in the *rrp4-G226D* cells.

### The Rrp4 M68T variant can associate with the RNA exosome complex

The sensitivity of the *rrp4-M68T* cells to drugs that impact RNA processing (Figure 3D) and the observed accumulation of key RNA exosome target RNAs (Figure 4) suggest that RNA exosome function may be impaired by the modeled multiple myeloma amino acid substitution. Previous studies suggest that SHRF-linked amino acid substitutions modeled in Rrp4 affect the RNA exosome function in part by disrupting complex integrity (Sterrett *et al*. 2021). To assess the impact of the modeled multiple myeloma amino acid substitution on the association of Rrp4 with other RNA exosome core subunits, we first assayed the protein level of Rrp4 M68T. We measured the steady-state level of the Myc-tagged Rrp4 M68T subunit when expressed as the sole copy of the Rrp4 protein in *rrp4∆* cells grown at either 30°C or 37°C (Figure 5A). Immunoblotting reveals that the steady-state level of Rrp4 M68T is comparable to wild-type Rrp4 at both temperatures tested. We next performed co-immunoprecipitations using *RRP45-TAP* cells that contain the endogenous, genomic *RRP4* gene and express a C-terminally tandem affinity purification (TAP)-tagged Rrp45 core subunit from the endogenous *RRP45* locus. We expressed Rrp4-Myc or Rrp4 M68T-Myc from plasmids in these *RRP45-TAP* cells. The Rrp45-TAP protein was immunoprecipitated and association of the Myc-tagged Rrp4 variants was assayed by immunoblotting (Figure 5B). Under these conditions in which an endogenous copy of *RRP4* is present, we can detect association of Rrp4 M68T-Myc with Rrp45-TAP at levels equal to that of Rrp4-Myc. As a comparative control, we also performed co-immunoprecipitations with *RRP45-TAP* cells expressing an exogenous Rrp4 G226D-Myc variant. Under these conditions with an endogenous copy of *RRP4* present, we cannot detect association of Rrp4 G226D-Myc with the TAP-tagged core subunit, supporting previous observations (Sterrett *et al*. 2021). Taken together, these data suggest that Rrp4 M68T is biochemically similar to a wild-type Rrp4 subunit, and the multiple myeloma amino acid substitution likely has no impact on RNA exosome complex integrity.

**Figure 5.**
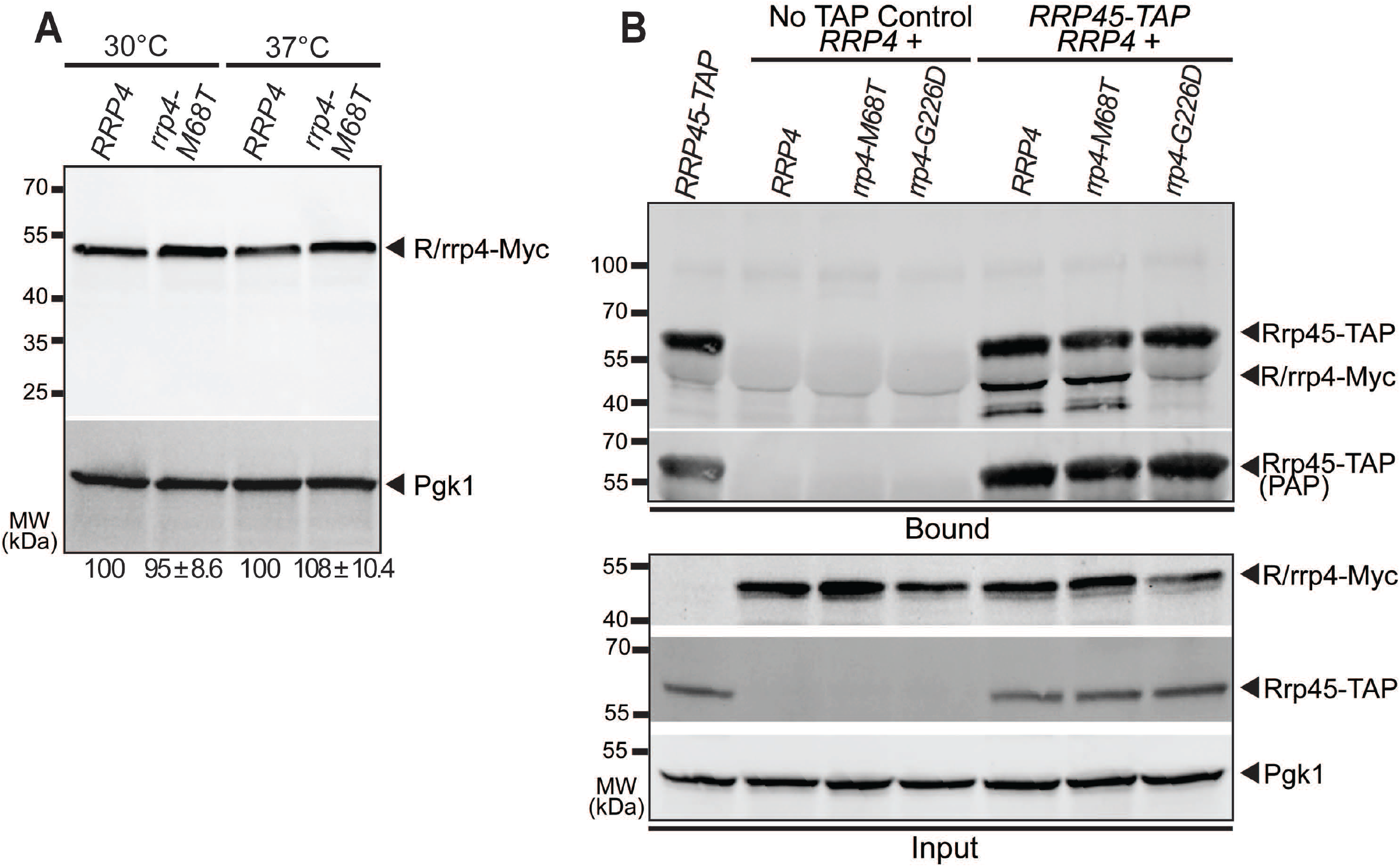
The modeled multiple myeloma amino acid substitution in Rrp4 does not impact Rrp4 protein level or association of the cap subunit with the RNA exosome complex. (A) The steady-state level of the Rrp4 M68T protein variant is equal to that of wild-type Rrp4 at both 30°C and 37°C. Lysates of *rrp4Δ* cells expressing Myc-tagged wild-type Rrp4 or Rrp4 M68T grown at 30°C or 37°C were analyzed by immunoblotting with an anti-Myc antibody. An anti-Pgk1 antibody was used to detect 3-phosphoglycerate kinase (Pgk1) as a loading control. The mean percentage of Rrp4 M68T-Myc normalized to Rrp4-Myc from four independent experiments (n = 4) is shown. Quantitation of the immunoblot was performed as described in *Materials and Methods*. (B) The Rrp4 M68T variant co-precipitates with the RNA exosome core subunit Rrp45 in the presence of a wild-type copy of Rrp4. Tandem affinity purification (TAP)-tagged Rrp45 was immunoprecipitated from *RRP45-TAP* cells expressing endogenous, wild-type Rrp4 and co-expressing Myc-tagged Rrp4, Rrp4 M68T, or, as a control, Rrp4 G226D grown at 30°C and bound (top) and input (bottom) samples were analyzed by immunoblotting. As a control, immunoprecipitations were also performed from untagged *RRP45* cells (No TAP Control) expressing Myc-tagged Rrp4 and Rrp4 variants. The bound/input level of Rrp4-Myc was detected with anti-Myc antibody and bound/input level of Rrp45-TAP was detected with a peroxidase anti-peroxidase (PAP) antibody. Bound Rrp45-TAP is also detected by the anti-Myc antibody as the Protein A moiety of the TAP tag binds to the antibody. The input level of 3-phosphoglycerate kinase (Pgk1) was detected with an anti-Pgk1 antibody and shown as a loading control. Data shown is representative of three independent experiments (n = 3).

### The *rrp4-M68T* mutant shows negative genetic interactions with *mtr4* mutants that impact TRAMP complex association and RNA helicase unwinding

As the Rrp4 M68T variant associates with the RNA exosome complex and has a steady level equivalent to wild-type Rrp4, the observed sensitivity to disrupted RNA processing in *rrp4-M68T* cells (Figure 3D) and accumulation of select RNA exosome target transcripts (Figure 4) could be due to altered interaction between Rrp4 and Mtr4. As depicted in Figure 6A, the nuclear RNA exosome cofactors Mpp6 and Rrp47 and the associated exonuclease Rrp6 aid in recruiting Mtr4 to the RNA exosome (Wasmuth *et al*. 2017). Rrp6 and Rrp47 form a composite site that binds to the N-terminus of Mtr4, recruiting the helicase to the RNA exosome (Schuch *et al*. 2014). Mtr4 forms contacts with the cap subunit Rrp4 and the cofactor Mpp6, stabilizing the helicase on the RNA exosome complex through a very conserved interface between the cap subunit and the helicase (Falk *et al*. 2017; Weick *et al*. 2018). The interaction between Mtr4 and Rrp4 provides a surface for the RNA exosome to associate with the TRAMP complex, which helps facilitates nuclear RNA surveillance and quality control of ncRNA (Lacava *et al*. 2005; Vaňáčová *et al*. 2005; Houseley and Tollervey 2008; Houseley and Tollervey 2009; Lubas *et al*. 2011; Stuparevic *et al*. 2013; Rodríguez-Galán *et al*. 2015; Kilchert *et al*. 2016b; Zinder and Lima 2017; Ogami *et al*. 2018; Belair *et al*. 2019). In addition to Mtr4, the TRAMP complex is composed of a zinc-knuckle RNA binding protein, Air1 or Air2, and a non-canonical oligo(A) polymerase, Trf4 or Tr45, that oligoadenylates RNA (Belair *et al*. 2018). The TRAMP complex triggers degradation by adding short polyadenylated tails to the 3’ end of substrate RNA and delivering them to the RNA exosome (Houseley *et al*. 2006; Anderson and Wang 2009; Belair *et al*. 2018; Ogami *et al*. 2018).

**Figure 6.**
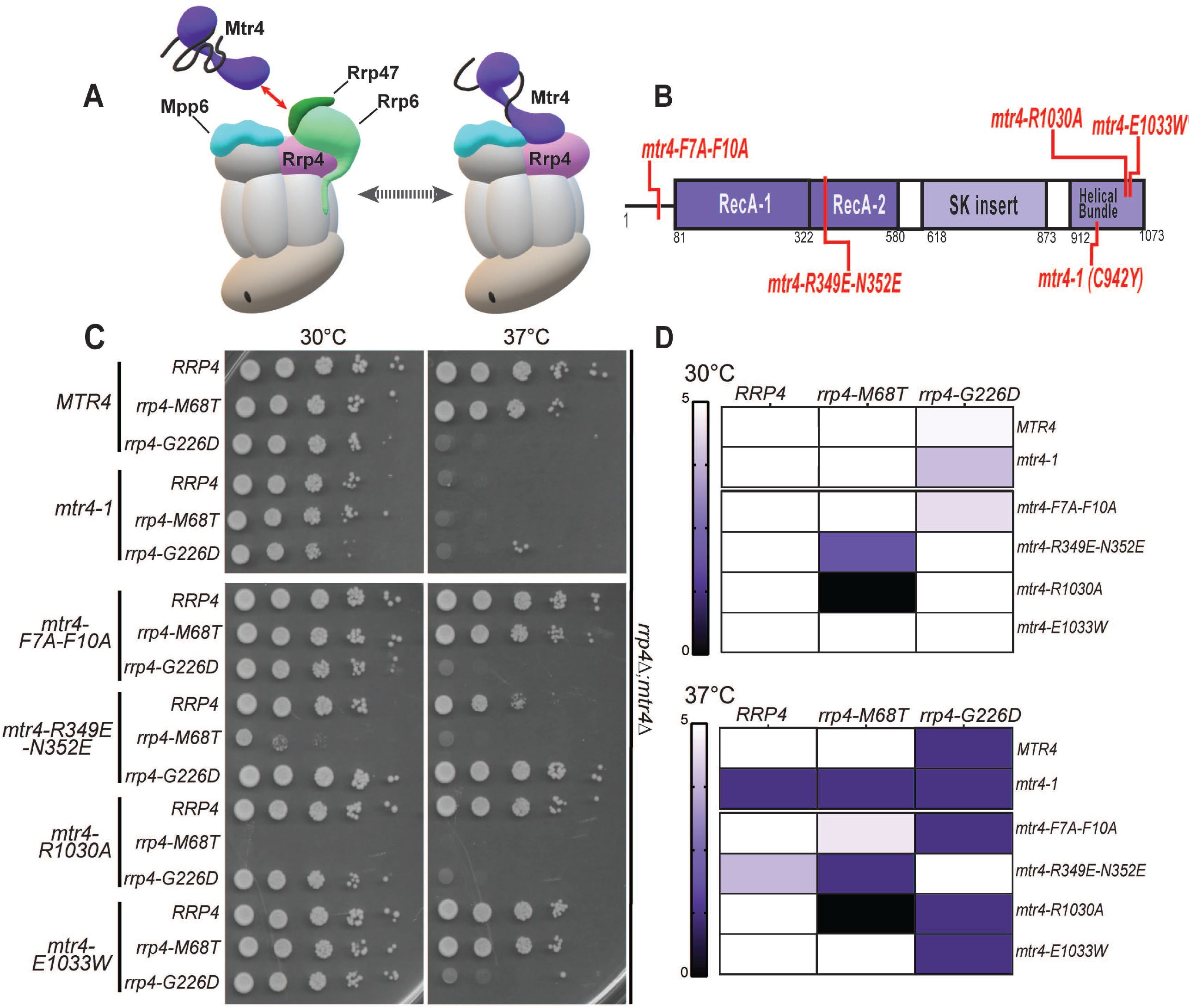
The *rrp4-M68T* mutant cells show specific negative genetic interactions with *mtr4* mutants that are predicted to impair the Trf4/5-Air1/2-Mtr4 (TRAMP) complex. (A) Cartoon depicting the budding yeast nuclear RNA exosome with interacting nuclear cofactors Mpp6 (turquoise) and Rrp47 (dark green), the exoribonuclease Rrp6 (light green), and the essential RNA helicase, Mtr4 (purple) (Schuch *et al*. 2014; Falk *et al*. 2017; Schuller *et al*. 2018). The association of Mtr4 with the RNA exosome is facilitated by interactions between Mtr4 and Rrp6/Rrp47 (denoted by the red arrowed line) and by interactions with Mpp6 which is associated with the Rrp40 RNA exosome subunit and the Rrp4 subunit (Weir *et al*. 2010; Schuch *et al*. 2014; Wasmuth *et al*. 2017). The association of Mtr4 with the RNA exosome can also facilitate interaction with the <u>Tr</u>f4/5-<u>A</u>ir1/2-<u>M</u>tr4 <u>p</u>olyadenylation (TRAMP) complex, which triggers degradation of certain RNA targets by adding short oligo(A) tails to the 3’ end of these targets and delivering them to the RNA exosome (Houseley *et al*. 2006; Anderson and Wang 2009; Belair *et al*. 2018; Ogami *et al*. 2018). In addition to Mtr4, the TRAMP complex is composed of a noncanonical poly(A) polymerase, Trf4/5, and a zinc-knuckle RNA binding protein, Air1/2 (Belair *et al*. 2018). Central to the degradation of TRAMP-targeted RNAs by the RNA exosome is the association of Mtr4 with Trf4/5, Air1/2 and the cap subunits and nuclear cofactors of the RNA exosome complex (Falk *et al*. 2014; Schuch *et al*. 2014). (B) Domain structure for *S. cerevisiae* Mtr4. The helicase has a low-complexity N-terminal sequence followed by the conserved helicase region. The helicase region is composed of two RecA domains and a helical domain (labeled helical bundle) that form the globular core typical of DExH family proteins. The helical bundle was originally described as the “ratchet” domain for its role in translocating nucleic acid by a Brownian ratchet (Büttner *et al*. 2007). In addition, Mtr4 contains an insertion domain and KOW domain that fold into a helical stalk (labeled SK insertion) (Jackson *et al*. 2010; Weir *et al*. 2010). The amino acid changes used for this experiment are labeled in red along the domain structure. (C) Double mutant cells containing *rrp4-M68T* and specific *mtr4* mutants show lethality at both 30°C and 37°C. The *rrp4∆ mtr4∆* double mutant cells were serially diluted, spotted onto solid media, and grown at the indicated temperatures for 3 days. The *mtr4* mutant plasmids included in this experiment are as follows; *mtr4-1—* a temperature sensitive mutant that contains a missense mutation resulting in the amino acid substitution Cys942Tyr, which causes accumulation of poly(A)+ RNA in the nucleus at 37°C (Kadowaki *et al*. 1994; Kadowaki *et al*. 1995; Liang *et al*. 1996); *mtr4-F7A-F10A—*an *mtr4* allele that impairs the interaction with Rrp6/Rrp47 (Schuch *et al*. 2014); *mtr4-R349E-N352E—*a mutation that impairs the association of Mtr4 with the poly(A) RNA polymerase Trf4 with the Mtr4 helicase (Falk *et al*. 2014); *mtr4R-1030A* and *mtr4-E1033W*—two mutations within the helical bundle that differentially impact nucleic acid unwinding by Mtr4 (Taylor *et al*. 2014). *mtr4-1* mutant cells expressing *RRP4, rrp4-M68T* or *rrp4-G226D* show lethality at 37°C presumably due to the known temperature sensitive nature of the *mtr4-1* allele (Liang *et al*. 1996). Growth of double mutant cells containing *rrp4-M68T* or *rrp4-G226D* are shown. (D) Summary of *rrp4 mtr4* mutant cell growth. Triplicate solid media assays were performed on double mutant cells containing *rrp4-M68T* or *rrp4-G226D* and the series of *mtr4* variants. Cell growth at both 30°C and 37°C was semi-quantified on a scale of 0 (lethal; black) to 5 (comparable to *RRP4* wild-type growth; white). Growth scale of the double mutant cells is represented through the color gradient on the two heatmaps. All double mutant cells were generated as described in *Materials and Methods*.

To assess genetic interactions between Mtr4 and the RNA exosome in *S. cerevisiae* modeling the multiple myeloma patient mutation, we performed an analysis of a series of five *mtr4* mutant alleles that introduce amino acid substitutions in Mtr4 as summarized in Figure 6B. We also included the *rrp4* mutant variant, *rrp4-G226D*, for comparison as this *rrp4* variant has a negative interaction with the *mtr4-F7A-F10A* mutant allele (Sterrett *et al*. 2021). Included within our five *mtr4* alleles is the *mtr4-1* allele which is a known temperature sensitive mutant (Kadowaki *et al*. 1994; Kadowaki *et al*. 1995; Liang *et al*. 1996; Weir *et al*. 2010). These genetic data are shown both as representative solid growth assays (Figure 6C) and as a heatmap (Figure 6D), which summarizes data from three independent replicates for all these genetic experiments. As predicted, *RRP4 mtr4-1* cells have a severe growth defect at 37°C that is shared by both double mutant *rrp4-M68T mtr4-1* and *rrp4-G226D mtr4-1*.

Two of the five *mtr4* mutant alleles, *mtr4-F7A-F10A* and *mtr4-R349E-N352E*, impair protein-protein interactions of Mtr4 in *S. cerevisiae*. The Mtr4 F7A F10A variant disrupts Mtr4 interactions with Rrp6/Rrp47 by introducing two amino acid substitutions, F7A and F10A, into the N-terminus of Mtr4 (Figure 6B) (Schuch *et al*. 2014). The *mtr4-R349E-N352E* mutant allele impairs the association of Mtr4 with the poly(A) RNA polymerase Trf4 within the TRAMP complex and thus disrupts the recruitment of the TRAMP complex to the RNA exosome (Falk *et al*. 2014). The *rrp4-M68T mtr4-F7A-F10A* double mutant cells grow similarly to *rrp4-M68T* cells at 30°C, however at 37°C the double mutant cells show a mild growth defect in comparison to the single mutant *rrp4-M68T* and the *RRP4 mtr4-F7A-F10A* cells (Figure 6D). As shown previously, the *rrp4-G226D mtr4-F7A-F10A* show a severe growth defect at 37°C compared to *rrp4-G226D* cells (Sterrett *et al*. 2021). The *rrp4 M68T mtr4-R349E-N352E* double mutant cells show severe growth defects at 30°C and 37°C and compared to the single mutant *rrp4-M68T* cells or the *RRP4 mtr4-R349E-N352E* control cells. Contrastingly, the *rrp4 G226D mtr4-R349E-N352E* double mutant cells show no difference in growth at 30°C compared to the *RRP4 mtr4-R349E-N352E* control cells and improved growth compared to the single mutant *rrp4-G226D* cells at 37°C (Figure 6C and 6D).

The final two *mtr4* mutant alleles that we tested for genetic interaction with the *rrp4-M68T* variant impact nucleic acid unwinding by Mtr4 (Taylor *et al*. 2014). Studies of an RNA-bound Mtr4 structure demonstrate that residues of R1030 and E1033 mediate key nucleic acid base interactions with the helicase helical bundle (Weir *et al*. 2010; Taylor *et al*. 2014). Mutagenesis of these residues in *S. cerevisiae*, generating the mutant alleles *mtr4-R1030A* and *mtr4-E1033A*, reveal that these residues play important but distinct roles in helicase activity (Taylor *et al*. 2014). The *rrp4-M68T mtr4-R1030A* double mutant cells are not viable at either temperature tested. In contrast the *rrp4-M68T mtr4-E1033A* double mutant cells are viable and grow similar to the *RRP4 mtr4-E1033A* control cells at both 30°C and 37°C. The *rrp4-G226D mtr4-R1030A* and *rrp4-G226D mtr4-E1033A* double mutant cells show lethality only at 37°C but have comparable growth to the single mutant *rrp4-G226D* cells as well as the *RRP4 mtr4-R1030A* and *RRP4 mtr4-E1033A* control cells at 30°C.

The *rrp4-M68T* double mutants that show synthetical lethality are viable when rescued by expression of a wild-type *RRP4* plasmid (Figure S3), demonstrating that the growth defects and lethality observed are due to negative genetic interactions between the *rrp4* and *mtr4* mutants. Taken together, these data suggest that the *rrp4-M68T* cells have negative genetic interactions with specific *mtr4* mutant alleles, distinct from those previously described for the *rrp4-G226D* mutant model.

### The *rrp4-M68T* mutant shows negative genetic interactions with *mpp6Δ*

As depicted in Figure 6A, the nuclear RNA exosome cofactors Mpp6 and Rrp47 and the exoribonuclease Rrp6 help to recruit and stabilize the interaction with Mtr4. The exosome cofactor Rrp47 interacts with and stabilizes the exoribonuclease Rrp6 and the cofactor Mpp6 interacts with the nuclear RNA exosome through direct contacts with the cap subunit Rrp40 (Schuch *et al*. 2014; Wasmuth *et al*. 2014; Wasmuth *et al*. 2017). To further evaluate the impact that the modeled multiple myeloma amino acid substitution may have on the RNA exosome-Mtr4 interaction *in vivo*, we tested whether the *rrp4-M68T* variant exhibits genetic interactions with *mpp6* or *rrp47* mutants by deleting these non-essential, nuclear exosome cofactor genes *MPP6* and *RRP47* in combination with *rrp4-M68T*. For comparison, we included the *rrp4-G226D* variant as these cells have known negative genetic interactions with these mutants (Sterrett *et al*. 2021). We examined the growth of these double mutants relative to single mutants (*rrp4-M68T* or *rrp4-G226D*) and the control mutant cells (*RRP4 mpp6Δ* or *RRP4 rrp47Δ*) in solid and liquid media growth assays (Figure 7). In the solid media growth assays the *rrp4-M68T mpp6Δ* double mutant cells show growth very similar to the *rrp4-M68T* cells at both 30°C and 37°C after both one and two days of growth (Figure 7B). The *rrp4-M68T rrp47Δ* cells show a severe growth defect at 37°C compared to the single mutant *rrp4-M68T;* however, this impaired growth is comparable to that of the *RRP4 rrp47Δ* cells, which has been previously reported for the single mutant *rrp47Δ* (Briggs *et al*. 1998; Mitchell *et al*. 2003) (Figure 6C). In contrast, the *rrp4-G226D mpp6Δ* and *rrp4-G226D rrp47Δ* double mutant cells show a severe growth defect at both temperatures compared to the single mutant *rrp4-G226D* cells as described previously (Sterrett *et al*. 2021).

**Figure 7.**
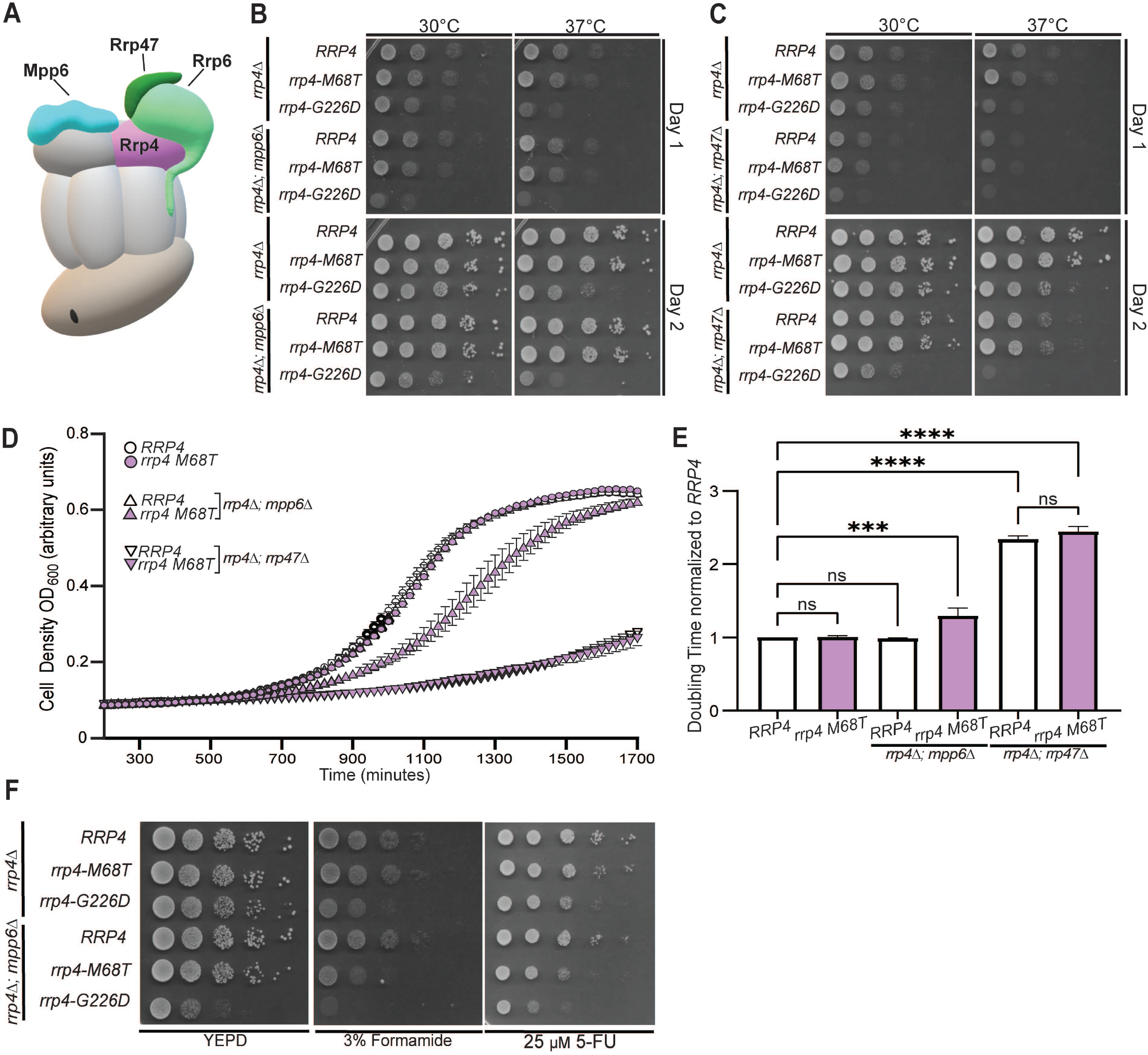
The *rrp4-M68T mpp6*Δ double mutant cells exhibits impaired growth that is exacerbated on drugs that impact RNA processing. (A) Cartoon schematic of the budding yeast nuclear RNA exosome in complex with nuclear cofactors Mpp6 (turquoise) and Rrp6/47 (light green/dark green) (Schuller *et al*. 2018). Serial dilution growth assays of double mutant (B) *rrp4-M68T mpp6∆* or (C) *rrp4-M68T rrp47∆* cells at 30°C and 37°C. The double mutant cells (*rrp4∆* with *mpp6∆*, or *rrp47∆)* containing control *RRP4* or *rrp4* variants *rrp4-M68T* and *rrp4-G226D* plasmids were serially diluted, spotted onto selective solid media, and grown at the indicated temperatures for two days. The double mutant cells *rrp4-G226D mpp6∆* and *rrp4-G226D rrp47∆* were included as a comparative control and show growth defects as described previously (Sterrett *et al*. 2021). Data shown are representative of three independent assays (n = 3). (D) and (E) Double mutant cells containing *rrp4-G226D* and *mpp6∆* exhibit a statistically significant increase in doubling time in liquid culture. Double mutant cells (*rrp4∆ mpp6∆* or *rrp4∆ rrp47∆)* containing control *RRP4* or *rrp4-M68T* plasmids were diluted in selective media and grown at 37°C with optical density measurements used to assess cell density over time. Data shown is collected from four independent samples (n = 4). (E) Doubling time for each sample was quantified and normalized to the growth rate of control *RRP4* cells. All calculations were performed as described in *Materials and Methods*. Full liquid growth curves of both *rrp4-M68T mpp6∆* and *rrp4-M68T rrp47∆* mutant cells are shown in Supplemental Figure S4. (F) Double mutant cells *rrp4-M68T mpp6∆* exhibit impaired growth on solid media containing drugs impacting RNA processing. The *rrp4∆ mpp6∆* cells expressing *RRP4, rrp4-M68T* or *rrp4-G226D* were serially diluted, spotted onto solid YEPD media containing 3% formamide or selective media containing 25 µM fluorouracil (5-FU) and grown at 30°C for three days. The *rrp4-M68T mpp6∆* cells show impaired growth when compared to *RRP4 mpp6∆* cells. The *rrp4-G226D mpp6∆* cells show exacerbated growth defects on 3% formamide and 25 µM 5-FU at 30°C. Data shown are representative of three independent assays (n = 3).

While the solid media growth assay suggests comparable growth between the controls and the *rrp4-M68T* double mutant cells, the liquid media growth assay reveals a modest growth defect of the *rrp4-M68T mpp6Δ* at 37°C compared to both the *rrp4-M68T* and control *RRP4 mpp6Δ* cells (Figure 7D), with the doubling time significantly longer than that of wild-type *RRP4* cells (Figure 7E). The liquid growth assay also shows doubling times for *rrp4-M68T rrp47Δ* and *RRP4 rrp47Δ* double mutants are nearly twice that of wild-type *RRP4* cells, but do not differ significantly when compared to each other (Figure 7D and 7E). The observed growth defect of the *rrp4-M68T mpp6Δ* double mutant in liquid culture is revealed in a solid media growth assay when the cells are challenged with formamide or 5-FU (Figure 7F). The distinct growth defect of the *rrp4-G226D mpp6Δ* double mutant is also exacerbated by growth on these chemicals. Taken together, these data suggest a negative genetic interaction between the *rrp4* variants and *mpp6* mutants, with both double mutants having exacerbated defects when challenged with drugs that impact RNA metabolism.

## DISCUSSION

In this study, we modeled and analyzed a multiple myeloma patient *EXOSC2* mutation in the *S. cerevisiae* homolog *RRP4*. We generated *rrp4-M68T* mutant cells expressing the variant Rrp4 M68T, which corresponds to the EXOSC2 M40T variant. Analysis of these *rrp4-M68T* cells reveals that this amino acid substitution affects RNA exosome function. While our biochemical assays show that the Rrp4 M68T variant can associate with the RNA exosome complex and function as the sole copy of the essential Rrp4 RNA exosome cap subunit, *rrp4-M68T* cells do show growth defects when grown in media containing drugs that impact RNA processing. The *rrp4-M68T* cells also show accumulation of known RNA exosome targets. These defects in RNA exosome function could result from an impaired interaction between the complex and the essential RNA helicase Mtr4 as predicted by structural modeling. Our genetic analyses support this model as *rrp4-M68T* cells show negative genetic interactions with both *mpp6* and *mtr4* mutants. These data suggest that the introduction of the multiple myeloma associated amino acid change could impact the binding interface between EXOSC2 and MTR4, potentially impairing the function of the essential RNA exosome *in vivo* for a subset of Mtr4-dependent targets.

Structural studies reveal the evolutionary conservation of the interaction between the RNA exosome and Mtr4 (Figure S1), with the helicase cofactor interacting with the complex through multiple points of contact including a direct interface with EXOSC2/Rrp4 and indirect stabilizing interactions with the cofactors Mpp6, Rrp47 and the associated exonuclease Rrp6 (Falk *et al*. 2017; Weick *et al*. 2018). This robust interaction between the complex and the essential helicase likely explains why the *rrp4-M68T* cells show no functional consequences unless challenged through introduction of drugs impacting RNA processing or loss of other stabilizing cofactors, such as in *rrp4-M68T mpp6Δ* double mutant cells. While this model would also predict a negative genetic interaction between *rrp47Δ* and *rrp4-M68T*, the *rrp4-M68T rrp47Δ* cells show a growth defect at 37°C that is indistinguishable from that of *RRP4 rrp47Δ* cells. This growth defect in *rrp4-M68T rrp47Δ* and *RRP4 rrp47Δ* cells is likely due to the loss of Rrp6 association with the RNA exosome complex given the stabilizing role Rrp47 plays for Rrp6 (Mitchell *et al*. 2003; Wasmuth *et al*. 2014). The growth defects resulting from destabilization of Mtr4 in *rrp4-M68T rrp47Δ* cells is likely masked by the larger consequence of disassociating Rrp6 from the complex. We do detect a slight growth defect in *rrp4-M68T* cells expressing an *mtr4* variant that disrupts the stabilizing interactions between Rrp6, Rrp47 and Mtr4 (*mtr4-F7A-F10A)*, pointing to the importance of the Rrp4-Mtr4 interface. Furthermore, we observe molecular consequences in *rrp4-M68T* cells as accumulation of several documented RNA exosome target transcripts. These data suggest that the consequences resulting *in vivo* from the Rrp4 M68T variant are subtle though could be impactful for a specific set of target RNAs by the RNA exosome complex.

The interaction between the RNA exosome and Mtr4 could also be critical for other interactions, particularly those involving the TRAMP (Trf4/5-Air1/2-Mtr4 Polyadenylation) complex. Our genetic analyses reveal a negative genetic interaction between *rrp4-M68T* and *mtr4-R349E-N352E*. The Mtr4 R349E N352E variant impairs Mtr4-Trf4 binding and impacts TRAMP complex assembly *in vivo* (Falk *et al*. 2014). The *rrp4-M68T* cells that express Mtr4 R349E N352E as the sole copy of the helicase grow very poorly at both 30°C and 37°C as compared to control *RRP4 mtr4-R349E-N352E* cells, suggesting TRAMP complex assembly and association with the RNA exosome may also be impacted by Rrp4 M68T. Intriguingly, we also detect synthetic lethality for the *rrp4-M68T mtr4-R1030A* double mutant. This lethality is specific to *rrp4-M68T mtr4-R1030A* cells as the *rrp4-M68T* cells expressing the other helicase mutant, *mtr4-E1033W*, show growth similar to the control (*RRP4 mtr4-E1033W)*. Both Mtr4 R1030A and Mtr4 E1033W decrease helicase unwinding capability (Taylor *et al*. 2014). However, the Mtr4 R1030A variant is also implicated in disrupting target discrimination by the TRAMP complex, potentially by disrupting preferential polyadenylation by Trf4 (Taylor *et al*. 2014). Therefore, the negative genetic interaction observed for *rrp4-M68T rrp4-R1030A* cells further suggests that TRAMP function is impacted in *rrp4-M68T* cells. Taken together with our structural modeling data, we hypothesize that a stabilized interaction between the RNA exosome and Mtr4 is necessary for TRAMP association and the slightest perturbation, even a subtle destabilization at one contact point with the helicase, could disrupt this vital interaction between TRAMP and the complex. More biochemical studies could be performed to explore how changes within the EXOSC2-Mtr4 interface impact the interaction with the TRAMP complex.

Our studies also show that *rrp4-M68T* mutant cells have distinct genetic interactions as compared to the *rrp4-G226D* cells. The *rrp4-G226D mtr4-R349E-N352E* double mutant cells surprisingly show improved growth at 37°C compared to either single mutant. Even more surprising is the synthetic lethality in cells expressing *rrp4-G226D* and either *mtr4* helicase mutant (*mtr4-R1030A* and *mtr4-E1033W*). These genetic interactions could suggest that the modeled SHRF amino acid substitution (Rrp4 G226D) has distinct *in vivo* consequences compared to the modeled multiple myeloma-associated substitution Rrp4 M68T. The Rrp4 G226D variant has decreased association with Mtr4 and the *rrp4-G226D* cells show transcriptomic differences from wild-type cells that suggest disrupted RNA exosome-Mtr4 interactions (Sterrett *et al*. 2021). Some RNA exosome targets also accumulate in *rrp4-M68T* cells that accumulate in *rrp4-G226D* cells. However notably we do not detect any changes in select CUTs or 5.8S rRNA precursors in *rrp4-M68T* cells. These data suggest that there may be some distinct defects in RNA exosome function in *rrp4-G226D* cells compared to the *rrp4-M68T* cells though they may also have some overlapping consequences *in vivo* due to altered association with Mtr4. The difference in molecular consequences between the two *rrp4* variants could be attributed to the impact on RNA exosome complex integrity observed in *rrp4-G226D* cells, which was not observed in *rrp4-M68T* cells (Figure 5B) (Sterrett *et al*. 2021).

The difference in severity of functional and molecular consequences we observe for the *rrp4-M68T* and *rrp4-G226D* mutant models may partially explain the differences in disease pathology between SHRF patients with the mutation *EXOSC2 G198D* and the multiple myeloma patient with the mutation *EXOSC2 M40T*. The *EXOSC2 G198D* mutation was identified in SHRF patients through whole exome sequencing and classified as causing a novel Mendelian syndrome (Di Donato *et al*. 2016). In contrast, the *EXOCS2 M40T* mutation is a spontaneous, somatic mutation that likely co-occurred with a chromosome 9 duplication. Additionally, the patient with this *EXOSC2 M40T* mutation has several chromosomal aberrations that are a hallmark of multiple myeloma, suggesting these *EXOSC2* mutations could be passenger mutations rather than a pathogenic driver of the multiple myeloma. Upon further analysis of the noncoding mutations in the patient harboring *EXOSC2 M40T*, we found a second mutation present in intron 1 of *EXOSC2* in this patient (Figure S5). This mutation *(EXOSC2 SNV chr9:130,693,915 T>G)* is predicted to alter the splice donor site and likely result in a misprocessed mRNA or truncated protein. Through RNA-Seq data available in CoMMpass for this patient, we determined that the *EXOSC2 M40T* missense mutation and the splice donor mutation are expressed from the same allele. Interestingly, we calculate the allelic frequency of these two *EXOSC2* mutations to be very similar (0.2266 vs. 0.2191). This suggests that *EXOSC2* M40T and the *EXOSC2* splice donor mutation either co-occurred or that the splice donor mutation was selected for in response to the *EXOSC2 M40T* missense mutation, which could negatively affect cell growth and/or survival. As *EXOCS2* is an essential gene in 1076/1086 cancer cell lines in the Cancer Dependencies Map project (depmap.org) including all 19 myeloma cell lines in the dataset, a future approach would be to CRISPR mutate the *EXOSC2* M40T mutation within myeloma cell lines to determine the effects on exosome function and myeloma cell growth and survival.

As the *rrp4-M68T* cells show defects in function of the RNA exosome likely through altered of interactions with the RNA helicase Mtr4 and the associated TRAMP complex, this EXOSC2 M40T substitution could be detrimental to the function of the human RNA exosome. Altering key cofactor interactions with the RNA exosome could impact the processing and degradation of target RNA transcripts such as small ncRNA species that have key regulatory roles in various cellular processes. Furthermore, the interaction between the RNA exosome and MTR4 has been suggested to resolve secondary DNA structures associated with strand asymmetric DNA mutagenesis that can lead to genome instability and chromosomal translocations particularly in plasma B cells (Lim *et al*. 2017). The high level of evolutionary conservation within the N-terminus of EXOSC2 that interacts with MTR4 (Figure 1C and Figure S1), suggests that there could be evolutionary pressure to maintain the integrity of certain sequences within EXOSC2 that specifically interact with key cofactors. Taking a genetic approach to assess different *EXOSC* missense mutations associated with human diseases can help unravel different consequences in specific interactions of the essential RNA exosome complex.

Utilizing the yeast genetic model system, we have characterized an *EXOSC2* mutation found in a multiple myeloma patient. However, this mutation was one of several mutations identified in genes encoding structural subunits of the RNA exosome in the CoMMpass study. The frequency of multiple myeloma mutations identified in the *DIS3* catalytic exosome gene suggests that there is an important link between multiple myeloma and the RNA exosome. By modeling identified *EXOSC* mutations in the budding yeast system, we can examine whether these mutations impair the function of the essential RNA exosome and provide a deeper understanding of the role this conserved complex may have in cancer pathologies. While it is unlikely that the identified *EXOSC* mutations drive the multiple myeloma disease, our study here clearly shows that these mutations have *in vivo* consequences for the conserved and essential RNA exosome-Mtr4 interaction. In addition, by studying other models of *EXOSC* disease linked mutations, such as those identified in RNA exosomopathy patients, we can provide insight into the biological pathways that are altered in these different disorders. As more pathogenic mutations are uncovered in *EXOSC* genes through patient genomic screenings, generating *in vivo* models to explore the consequences of these changes can help to define the most critical interactions of the complex with various cofactors thus expand our understanding of the biological functions of this singular, essential RNA processing and degradation complex.

## ACKNOWLEDGEMENTS

We thank members of the Corbett lab for critical discussions and input. We thank Dr. Benjamin Barwick for his contributions in assisting with the analysis of CoMMpass dataset. We also thank Dr. Graeme Conn and Dr. Ambro van Hoof for their intellectual contributions. This work was supported by National Institutes of Health (NIH) R01 grants (GM058728) and the NIH-funded Emory Initiative for Maximizing Student Development (R25 GM099644) to A.H.C and National Cancer Institute (NCI) R21 grant (CA273773) to L.H.B. M.C.S. was supported by a National Institute of General Medical Sciences (NIGMS) F31 grant (GM134649-01). S.E.S. was supported by the National Science Foundation (NSF) Graduate Research Fellowship (GRFP 1937971). Lastly, we would also like to thank the Genetics Society of America and the <u>Saccharomyces Genomic Database</u> (SGD) (Cherry *et al*. 2012) for providing community resources and support for scientific discovery.

## REFERENCES

Adams, A., D. E. Gottschling, C. A. Kaiser and T. Stearns, 1997 Methods in Yeast Genetics. Cold Spring Harbor Laboratory Press, Cold Spring Harbor.

Alexander, D. D., P. J. Mink, H. O. Adami, P. Cole, J. S. Mandel et al., 2007 Multiple myeloma: a review of the epidemiologic literature. Int J Cancer 120 Suppl 12: 40–61.

Allmang, C., J. Kufel, G. Chanfreau, P. Mitchell, E. Petfalski et al., 1999 Functions of the exosome in rRNA, snoRNA and snRNA synthesis. Embo Journal 18: 5399–5410.

Anderson, J. T., and X. Wang, 2009 Nuclear RNA surveillance: no sign of substrates tailing off. Crit Rev Biochem Mol Biol 44: 16–24.

Ashkenazy, H., S. Abadi, E. Martz, O. Chay, I. Mayrose et al., 2016 ConSurf 2016: an improved methodology to estimate and visualize evolutionary conservation in macromolecules. Nucleic Acids Research 44: W344–W350.

Ashkenazy, H., E. Erez, E. Martz, T. Pupko and N. Ben-Tal, 2010 ConSurf 2010: calculating evolutionary conservation in sequence and structure of proteins and nucleic acids. Nucleic Acids Research 38: W529–W533.

Barwick, B. G., P. Neri, N. J. Bahlis, A. K. Nooka, M. V. Dhodapkar et al., 2019 Multiple myeloma immunoglobulin lambda translocations portend poor prognosis. Nature Communications 10: 1911.

Belair, C., S. Sim, K. Y. Kim, Y. Tanaka, I. H. Park et al., 2019 The RNA exosome nuclease complex regulates human embryonic stem cell differentiation. J Cell Biol.

Belair, C., S. Sim and S. L. Wolin, 2018 Noncoding RNA Surveillance: The Ends Justify the Means. Chem Rev 118: 4422–4447.

Bergsagel, P. L., E. Nardini, L. Brents, M. Chesi and W. M. Kuehl, 1997 IgH translocations in multiple myeloma: a nearly universal event that rarely involves c-myc. Curr Top Microbiol Immunol 224: 283–287.

Boczonadi, V., J. S. Mueller, A. Pyle, J. Munkley, T. Dor et al., 2014 EXOSC8 mutations alter mRNA metabolism and cause hypomyelination with spinal muscular atrophy and cerebellar hypoplasia. Nature Communications 5.

Boyle, E. M., C. Ashby, R. G. Tytarenko, S. Deshpande, H. Wang et al., 2020 BRAF and DIS3 Mutations Associate with Adverse Outcome in a Long-term Follow-up of Patients with Multiple Myeloma. Clin Cancer Res 26: 2422–2432.

Briggs, M. W., K. T. Burkard and J. S. Butler, 1998 Rrp6p, the yeast homologue of the human PM-Scl 100-kDa autoantigen, is essential for efficient 5.8 S rRNA 3’ end formation. J Biol Chem 273: 13255–13263.

Burns, D., D. Donkervoort, D. Bharucha-Goebel, M. Giunta, B. Munro et al., 2017 A recessive mutation in EXOSC9 causes abnormal RNA metabolism resulting in a novel form of cerebellar hypoplasia/atrophy with early motor neuronopathy. Neuromuscul Disord 27: S38.

Burns, D. T., S. Donkervoort, J. S. Muller, E. Knierim, D. Bharucha-Goebel et al., 2018 Variants in EXOSC9 Disrupt the RNA Exosome and Result in Cerebellar Atrophy with Spinal Motor Neuronopathy. Am J Hum Genet 102: 858–873.

Büttner, K., S. Nehring and K. P. Hopfner, 2007 Structural basis for DNA duplex separation by a superfamily-2 helicase. Nat Struct Mol Biol 14: 647–652.

Celniker, G., G. Nimrod, H. Ashkenazy, F. Glaser, E. Martz et al., 2013 ConSurf: Using Evolutionary Data to Raise Testable Hypotheses about Protein Function. Israel Journal of Chemistry 53: 199–206.

Cervelli, T., and A. Galli, 2021 Yeast as a Tool to Understand the Significance of Human Disease-Associated Gene Variants. Genes (Basel) 12.

Chapman, M. A., M. S. Lawrence, J. J. Keats, K. Cibulskis, C. Sougnez et al., 2011 Initial genome sequencing and analysis of multiple myeloma. Nature 471: 467–472.

Cherry, J. M., E. L. Hong, C. Amundsen, R. Balakrishnan, G. Binkley et al., 2012 Saccharomyces Genome Database: the genomics resource of budding yeast. Nucleic Acids Res 40: D700–705.

Coy, S., A. Volanakis, S. Shah and L. Vasiljeva, 2013 The Sm complex is required for the processing of non-coding RNAs by the exosome. PloS one 8: e65606–e65606.

Da, B., D. Dawson and T. Stearns, 2000 Methods In Yeast Genetics: A Cold Spring Harbor Laboratory Course Manual.

de Amorim, J., Slavotinek, A., Fasken, M.B., Corbett, A.H., Morton, D.J, 2020 Modeling Pathogenic Variants in the RNA Exosome. RNA & Disease 7.

de la Cruz, J., D. Kressler, D. Tollervey and P. Linder, 1998 Dob1p (Mtr4p) is a putative ATP-dependent RNA helicase required for the 3’ end formation of 5.8S rRNA in Saccharomyces cerevisiae. Embo j 17: 1128–1140.

Delan-Forino, C., C. Schneider and D. Tollervey, 2017 Transcriptome-wide analysis of alternative routes for RNA substrates into the exosome complex. PLOS Genetics 13: e1006699.

Di Donato, N., T. Neuhann, A.-K. Kahlert, B. Klink, K. Hackmann et al., 2016 Mutations in EXOSC2 are associated with a novel syndrome characterised by retinitis pigmentosa, progressive hearing loss, premature ageing, short stature, mild intellectual disability and distinctive gestalt. Journal of Medical Genetics 53: 419–425.

Donahue, T. F., and S. A. Henry, 1981 myo-Inositol-1-phosphate synthase. Characteristics of the enzyme and identification of its structural gene in yeast. J Biol Chem 256: 7077–7085.

Eggens, V. R. C., P. G. Barth, J.-M. F. Niermeijer, J. N. Berg, N. Darin et al., 2014 EXOSC3 mutations in pontocerebellar hypoplasia type 1: novel mutations and genotype-phenotype correlations. Orphanet Journal of Rare Diseases 9: 23.

Falk, S., F. Bonneau, J. Ebert, A. Kogel and E. Conti, 2017 Mpp6 Incorporation in the Nuclear Exosome Contributes to RNA Channeling through the Mtr4 Helicase. Cell Reports 20: 2279–2286.

Falk, S., J. R. Weir, J. Hentschel, P. Reichelt, F. Bonneau et al., 2014 The molecular architecture of the TRAMP complex reveals the organization and interplay of its two catalytic activities. Mol Cell 55: 856–867.

Fasken, M. B., J. S. Losh, S. W. Leung, S. Brutus, B. Avin et al., 2017 Insight into the RNA Exosome Complex Through Modeling Pontocerebellar Hypoplasia Type 1b Disease Mutations in Yeast. Genetics 205: 221-+.

Fasken, M. B., D. J. Morton, E. G. Kuiper, S. K. Jones, S. W. Leung et al., 2020 The RNA Exosome and Human Disease. Methods Mol Biol 2062: 3–33.

Ghaemmaghami, S., W. K. Huh, K. Bower, R. W. Howson, A. Belle et al., 2003 Global analysis of protein expression in yeast. Nature 425: 737–741.

Giaever, G., P. Flaherty, J. Kumm, M. Proctor, C. Nislow et al., 2004 Chemogenomic profiling: identifying the functional interactions of small molecules in yeast. Proc Natl Acad Sci U S A 101: 793–798.

Gillespie, A., J. Gabunilas, J. C. Jen and G. F. Chanfreau, 2017 Mutations of EXOSC3/Rrp40p associated with neurological diseases impact ribosomal RNA processing functions of the exosome in S. cerevisiae. RNA 23: 466–472.

Hoskins, J., and J. Scott Butler, 2007 Evidence for distinct DNA- and RNA-based mechanisms of 5-fluorouracil cytotoxicity in Saccharomyces cerevisiae. Yeast 24: 861–870.

Hou, D., M. Ruiz and E. D. Andrulis, 2012 The ribonuclease Dis3 is an essential regulator of the developmental transcriptome. Bmc Genomics 13.

Houseley, J., J. LaCava and D. Tollervey, 2006 RNA-quality control by the exosome. Nat Rev Mol Cell Biol 7: 529–539.

Houseley, J., and D. Tollervey, 2008 The nuclear RNA surveillance machinery: the link between ncRNAs and genome structure in budding yeast? Biochim Biophys Acta 1779: 239–246.

Houseley, J., and D. Tollervey, 2009 The many pathways of RNA degradation. Cell 136: 763–776.

Hoyos-Manchado, R., F. Reyes-Martín, C. Rallis, E. Gamero-Estévez, P. Rodríguez-Gómez et al., 2017 RNA metabolism is the primary target of formamide in vivo. Scientific Reports 7: 15895.

Jackson, R. N., A. A. Klauer, B. J. Hintze, H. Robinson, A. van Hoof et al., 2010 The crystal structure of Mtr4 reveals a novel arch domain required for rRNA processing. EMBO J 29: 2205–2216.

Kadowaki, T., S. Chen, M. Hitomi, E. Jacobs, C. Kumagai et al., 1994 Isolation and characterization of Saccharomyces cerevisiae mRNA transport-defective (mtr) mutants. J Cell Biol 126: 649–659.

Kadowaki, T., R. Schneiter, M. Hitomi and A. M. Tartakoff, 1995 Mutations in nucleolar proteins lead to nucleolar accumulation of polyA+ RNA in Saccharomyces cerevisiae. Mol Biol Cell 6: 1103–1110.

Kilchert, C., S. Wittmann and L. Vasiljeva, 2016a The regulation and functions of the nuclear RNA exosome complex. Nature Reviews Molecular Cell Biology 17: 227–239.

Kilchert, C., S. Wittmann and L. Vasiljeva, 2016b The regulation and functions of the nuclear RNA exosome complex. Nat Rev Mol Cell Biol 17: 227–239.

Klig, L. S., and S. A. Henry, 1984 Isolation of the yeast INO1 gene: located on an autonomously replicating plasmid, the gene is fully regulated. Proc Natl Acad Sci U S A 81: 3816–3820.

LaCava, J., J. Houseley, C. Saveanu, E. Petfalski, E. Thompson et al., 2005 RNA degradation by the exosome is promoted by a nuclear polyade nylation complex. Cell 121: 713–724.

Laubach, J., P. Richardson and K. Anderson, 2011 Multiple myeloma. Annu Rev Med 62: 249–264.

Liang, S., M. Hitomi, Y. H. Hu, Y. Liu and A. M. Tartakoff, 1996 A DEAD-box-family protein is required for nucleocytoplasmic transport of yeast mRNA. Mol Cell Biol 16: 5139–5146.

Lim, J., P. K. Giri, D. Kazadi, B. Laffleur, W. Zhang et al., 2017 Nuclear Proximity of Mtr4 to RNA Exosome Restricts DNA Mutational Asymmetry. Cell 169: 523–537 e515.

Lim, S. J., P. J. Boyle, M. Chinen, R. K. Dale and E. P. Lei, 2013 Genome-wide localization of exosome components to active promoters and chromatin insulators in Drosophila. Nucleic Acids Res 41: 2963–2980.

Livak, K. J., and T. D. Schmittgen, 2001 Analysis of relative gene expression data using real-time quantitative PCR and the 2(-Delta Delta C(T)) Method. Methods 25: 402–408.

Lohr, J. G., P. Stojanov, S. L. Carter, P. Cruz-Gordillo, M. S. Lawrence et al., 2014 Widespread genetic heterogeneity in multiple myeloma: implications for targeted therapy. Cancer Cell 25: 91–101.

Lorentzen, E., A. Dziembowski, D. Lindner, B. Seraphin and E. Conti, 2007 RNA channelling by the archaeal exosome. Embo Reports 8: 470–476.

Losh, J., 2018 Identifying Subunit Organization and Function of the Nuclear RNA Exosome Machinery, pp. University of Texas, UT GSBS Dissertations and Theses.

Lubas, M., M. S. Christensen, M. S. Kristiansen, M. Domanski, L. G. Falkenby et al., 2011 Interaction profiling identifies the human nuclear exosome targeting complex. Mol Cell 43: 624–637.

Makino, D. L., M. Baumgaertner and E. Conti, 2013 Crystal structure of an RNA-bound 11-subunit eukaryotic exosome complex. Nature 495: 70–75.

Milligan, L., L. Decourty, C. Saveanu, J. Rappsilber, H. Ceulemans et al., 2008 A yeast exosome cofactor, Mpp6, functions in RNA surveillance and in the degradation of noncoding RNA transcripts. Mol Cell Biol 28: 5446–5457.

Mitchell, P., E. Petfalski, R. Houalla, A. Podtelejnikov, M. Mann et al., 2003 Rrp47p is an exosome-associated protein required for the 3’ processing of stable RNAs. Mol Cell Biol 23: 6982–6992.

Mitchell, P., E. Petfalski, A. Shevchenko, M. Mann and D. Tollervey, 1997 The exosome: A conserved eukaryotic RNA processing complex containing multiple 3’->5’ exoribonucleases. Cell 91: 457–466.

Mitchell, P., E. Petfalski and D. Tollervey, 1996 The 3’ end of yeast 5.8S rRNA is generated by an exonuclease processing mechanism. Genes Dev 10: 502–513.

Moore, M. J., and N. J. Proudfoot, 2009 Pre-mRNA processing reaches back to transcription and ahead to translation. Cell 136: 688–700.

Morton, D. J., B. Jalloh, L. Kim, I. Kremsky, R. J. Nair et al., 2020 A Drosophila model of Pontocerebellar Hypoplasia reveals a critical role for the RNA exosome in neurons. PLoS Genet 16: e1008901.

Oddone, A., E. Lorentzen, J. Basquin, A. Gasch, V. Rybin et al., 2007 Structural and biochemical characterization of the yeast exosome component Rrp40. EMBO reports 8: 63–69.

Ogami, K., Y. Chen and J. L. Manley, 2018 RNA surveillance by the nuclear RNA exosome: mechanisms and significance. Noncoding RNA 4.

Parker, R., 2012 RNA Degradation in Saccharomyces cerevisae. Genetics 191: 671–702.

Pefanis, E., J. Wang, G. Rothschild, J. Lim, J. Chao et al., 2014 Noncoding RNA transcription targets AID to divergently transcribed loci in B cells. Nature 514: 389–393.

Preker, P., J. Nielsen, S. Kammler, S. Lykke-Andersen, M. S. Christensen et al., 2008 RNA Exosome Depletion Reveals Transcription Upstream of Active Human Promoters. Science 322: 1851–1854.

Rodrigues, C. H. M., Y. Myung, D. E. V. Pires and D. B. Ascher, 2019 mCSM-PPI2: predicting the effects of mutations on protein-protein interactions. Nucleic Acids Res 47: W338–w344.

Rodríguez-Galán, O., J. J. García-Gómez, D. Kressler and J. de la Cruz, 2015 Immature large ribosomal subunits containing the 7S pre-rRNA can engage in translation in Saccharomyces cerevisiae. RNA biology 12: 838–846.

Sambrook, J., E. F. Fritsch and T. Maniatis, 1989 Molecular Cloning: A Laboratory Manual. Cold Spring Harbor Laboratory Press, Cold Spring Harbor, New York.

Schaeffer, D., B. Tsanova, A. Barbas, F. P. Reis, E. G. Dastidar et al., 2009 The exosome contains domains with specific endoribonuclease, exoribonuclease and cytoplasmic mRNA decay activities. Nat Struct Mol Biol 16: 56–62.

Schneider, C., G. Kudla, W. Wlotzka, A. Tuck and D. Tollervey, 2012 Transcriptome-wide Analysis of Exosome Targets. Molecular Cell 48: 422–433.

Schneider, C., and D. Tollervey, 2013 Threading the barrel of the RNA exosome. Trends in Biochemical Sciences 38: 485–493.

Schuch, B., M. Feigenbutz, D. L. Makino, S. Falk, C. Basquin et al., 2014 The exosome-binding factors Rrp6 and Rrp47 form a composite surface for recruiting the Mtr4 helicase. Embo Journal 33: 2829–2846.

Schuller Jan, M., S. Falk, L. Fromm, E. Hurt and E. Conti, 2018 Structure of the nuclear exosome captured on a maturing preribosome. Science 360: 219–222.

Schuller, J. M., S. Falk, L. Fromm, E. Hurt and E. Conti, 2018 Structure of the nuclear exosome captured on a maturing preribosome. Science 360: 219–222.

Sikorski, R. S., and P. Hieter, 1989 A system of shuttle vectors and yeast host strains designed for efficient manipulation of DNA in Saccharomyces cerevisiae. Genetics 122: 19–27.

Slater, M. L., 1973 Effect of reversible inhibition of deoxyribonucleic acid synthesis on the yeast cell cycle. J Bacteriol 113: 263–270.

Slavotinek, A., D. Misceo, S. Htun, L. Mathisen, E. Frengen et al., 2020 Biallelic variants in the RNA exosome gene EXOSC5 are associated with developmental delays, short stature, cerebellar hypoplasia and motor weakness. Hum Mol Genet 29: 2218–2239.

Somashekar, P. H., P. Kaur, J. Stephen, V. S. Guleria, R. Kadavigere et al., 2021 Bi-allelic missense variant, p.Ser35Leu in EXOSC1 is associated with pontocerebellar hypoplasia. Clin Genet 99: 594–600.

Staals, R. H., A. W. Bronkhorst, G. Schilders, S. Slomovic, G. Schuster et al., 2010 Dis3-like 1: a novel exoribonuclease associated with the human exosome. Embo j 29: 2358–2367.

Sterrett, M. C., L. Enyenihi, S. W. Leung, L. Hess, S. E. Strassler et al., 2021 A budding yeast model for human disease mutations in the EXOSC2 cap subunit of the RNA exosome complex. RNA 27: 1046–1067.

Stuparevic, I., C. Mosrin-Huaman, N. Hervouet-Coste, M. Remenaric and A. R. Rahmouni, 2013 Cotranscriptional Recruitment of RNA Exosome Cofactors Rrp47p and Mpp6p and Two Distinct Trf-Air-Mtr4 Polyadenylation (TRAMP) Complexes Assists the Exonuclease Rrp6p in the Targeting and Degradation of an Aberrant Messenger Ribonucleoprotein Particle (mRNP) in Yeast. Journal of Biological Chemistry 288: 31816–31829.

Suzuki, H., K. Nagai, E. Akutsu, H. Yamaki and N. Tanaka, 1970 On the mechanism of action of bleomycin. Strand scission of DNA caused by bleomycin and its binding to DNA in vitro. J Antibiot (Tokyo) 23: 473–480.

Taylor, L. L., R. N. Jackson, M. Rexhepaj, A. K. King, L. K. Lott et al., 2014 The Mtr4 ratchet helix and arch domain both function to promote RNA unwinding. Nucleic acids research 42: 13861–13872.

Tomecki, R., K. Drazkowska, I. Kucinski, K. Stodus, R. J. Szczesny et al., 2014 Multiple myeloma-associated hDIS3 mutations cause perturbations in cellular RNA metabolism and suggest hDIS3 PIN domain as a potential drug target. Nucleic Acids Res 42: 1270–1290.

van Hoof, A., P. Lennertz and R. Parker, 2000 Yeast exosome mutants accumulate 3’-extended polyadenylated forms of U4 small nuclear RNA and small nucleolar RNAs. Molecular and cellular biology 20: 441–452.

Vaňáčová, Š., J. Wolf, G. Martin, D. Blank, S. Dettwiler et al., 2005 A New Yeast Poly(A) Polymerase Complex Involved in RNA Quality Control. PLOS Biology 3: e189.

Wan, J., M. Yourshaw, H. Mamsa, S. Rudnik-Schöneborn, M. P. Menezes et al., 2012 Mutations in the RNA exosome component gene EXOSC3 cause pontocerebellar hypoplasia and spinal motor neuron degeneration. Nat Genet 44: 704–708.

Wasmuth, E. V., K. Januszyk and C. D. Lima, 2014 Structure of an Rrp6-RNA exosome complex bound to poly(A) RNA. Nature 511: 435–439.

Wasmuth, E. V., J. C. Zinder, D. Zattas, M. Das and C. D. Lima, 2017 Structure and reconstitution of yeast Mpp6-nuclear exosome complexes reveals that Mpp6 stimulates RNA decay and recruits the Mtr4 helicase. Elife 6: 24.

Weick, E. M., M. R. Puno, K. Januszyk, J. C. Zinder, M. A. DiMattia et al., 2018 Helicase-Dependent RNA Decay Illuminated by a Cryo-EM Structure of a Human Nuclear RNA Exosome-MTR4 Complex. Cell 173: 1663–1677 e1621.

Weir, J. R., F. Bonneau, J. Hentschel and E. Conti, 2010 Structural analysis reveals the characteristic features of Mtr4, a DExH helicase involved in nuclear RNA processing and surveillance. Proc Natl Acad Sci U S A 107: 12139–12144.

Weissbach, S., C. Langer, B. Puppe, T. Nedeva, E. Bach et al., 2015 The molecular spectrum and clinical impact of DIS3 mutations in multiple myeloma. Br J Haematol 169: 57–70.

Wyers, F., M. Rougemaille, G. Badis, J.-C. Rousselle, M.-E. Dufour et al., 2005 Cryptic Pol II Transcripts Are Degraded by a Nuclear Quality Control Pathway Involving a New Poly(A) Polymerase. Cell 121: 725–737.

Yang, X., V. Bayat, N. DiDonato, Y. Zhao, B. Zarnegar et al., 2019 Genetic and genomic studies of pathogenic EXOSC2 mutations in the newly described disease SHRF implicate the autophagy pathway in disease pathogenesis. Human Molecular Genetics 29: 541–553.

Zinder, J. C., and C. D. Lima, 2017 Targeting RNA for processing or destruction by the eukaryotic RNA exosome and its cofactors. Genes & Development 31: 88–100.

